# PARP1 regulates the genomic ribonucleotide processing activity of TOP1 to prevent the formation of toxic TOP1-DNA adducts and the associated mutations

**DOI:** 10.64898/2026.04.16.719024

**Authors:** Edward James Sarrain, Quan Wang, Angel Caitlin Bondoy, Feng Guo, Qi Cao, Hengyao Niu

## Abstract

Ribonucleotides are frequently incorporated into the genome during DNA replication and are normally removed by RNase H2. Alternatively, DNA topoisomerase 1 (TOP1) can process genomic ribonucleotides through two sequential cleavage events, generating a 2′,3′-cyclic phosphate-terminated nick and, subsequently, a TOP1 cleavage complex (TOP1cc). In addition, TOP1cc formation can lead to 2- to 5-bp slippage deletions, recently identified as the ID4 cancer signature. Here, we show that human TOP1 is intrinsically mutagenic, with a tendency to undergo secondary cleavage and form TOP1 cleavage complexes. We further demonstrate that PARP1, a central single-strand break repair factor, suppresses TOP1 secondary cleavage, TOP1cc formation, and the resulting slippage mutations. This regulation requires PARP1 to interact with both TOP1 and the 2′,3′-cyclic phosphate-terminated nick produced by the initial TOP1 cleavage. Notably, a PARP1 mutant defective in TOP1 interaction is partially impaired in restoring PARP inhibitor sensitivity in PARP1-knockout cells, suggesting that PARP inhibitor cytotoxicity may partly result from PARP1 trapping at 2′,3′-cyclic phosphate-terminated nicks and blocking their processing by TOP1 and other factors. Together, these findings reveal a novel role for PARP1 in regulating TOP1-dependent ribonucleotide processing, thereby suppressing cancer-signature mutations and contributing to the mechanism of PARP inhibitor cytotoxicity.

## Introduction

Ribonucleotides (rNMPs) are frequently incorporated into the genome by the major replicative polymerases. In mice, about one million ribonucleotides are incorporated into the genome per cell cycle, with human cells estimated to have an even higher accumulation (around 3 million ribonucleotides per genome).^1,2^ The removal of single mis-incorporated ribonucleotides occurs mainly through the error-free ribonucleotide excision repair pathway (RER), which involves cleavage at the 5’ of the embedded ribonucleotide by RNase H2 followed by strand displacement and DNA flap removal.^3,4^ In humans, partial loss-of-function mutations in RNase H2 lead to a 10 to 25-fold increase in the abundance of genomic ribonucleotides.^5^ Mutations in RNase H2-associated genes are linked to two types of cancers, chronic lymphocytic leukemia and metastatic castration-resistant prostate cancer, as well as autoimmune diseases, including Aicardi-Goutières syndrome and systemic lupus erythematosus.^6^ Although the cancer pathology in RNase H2 mutants is understudied, the autoimmune pathology is believed to be the result of genomic instability in the absence of functional RNase H2 that subsequently leads to the activation of an innate immune response.^1,7^ One of the potential sources for such genomic instability is the removal of genomic ribonucleotides by topoisomerase 1 (TOP1).

Biochemically, it has been long known that eukaryotic TOP1, including human TOP1, can cleave at the site of a single mis-incorporated ribonucleotide and generate a nick terminated by a 5’-OH and a 2’, 3’-cyclic phosphate (CP).^8^ Next, TOP1 can act at a CP nick in two ways. Besides a direct ligation that restores the embedded ribonucleotide, genetic studies from budding yeast indicate that TOP1 can initiate an alternative pathway to remove the ribonucleotide.^9–11^ In this pathway, TOP1 removes the terminal CP via a secondary cleavage ∼2-5 bp upstream of the ribonucleotide, which releases the CP-containing segment, but generates a trapped TOP1 cleavage complex (TOP1cc) due to the absence of a proximal end to ligate with. In DNA regions harboring short tandem repeats, the TOP1cc from the secondary cleavage can catalyze ligation across the gap following DNA misalignment, which leads to the formation of slippage mutations.^9^ Notably, such slippage mutations have been recently identified as the ID4 cancer signature, with a frequency increase in human RNASEH2A knockout cells, RNase H2A being the catalytic subunit of the RNase H2 trimeric complex (together with RNase H2B and H2C).^12,13^

While the TOP1-derived CP nick repair is well characterized in yeast, a major difference between single-strand break (SSB) repair in yeast and humans is that human cells express poly(ADP-ribose) polymerase 1 (PARP1), a universal SSB repair factor known for its ability to sense and signal breaks in DNA. PARP1 is a nuclear enzyme that is activated by DNA breaks and catalyzes the poly(ADP-ribosyl)ation (PARylation) of itself and other substrates to recruit and regulate different repair proteins.^14–16^ PARP inhibitors have become key agents in cancer therapy, where their efficacy arises not only from blocking PARP1’s catalytic use of NAD⁺ ligand but also from stabilizing PARP1-DNA complexes.^17,18^ This process, known as PARP trapping, enhances cytotoxicity in homologous recombination (HR)-deficient tumors, such as BRCA-deficient breast and ovarian cancers.^19–21^ Notably, RER-deficient cells are also hypersensitive to PARP inhibitor treatments, linking the processing of genomic ribonucleotides by TOP1 to PARP-trapping lesions.^22^ This association prompted us to investigate genomic ribonucleotide excision by human TOP1 and its regulation by PARP1. Our findings reveal the mutagenic nature of human TOP1 and how PARP1 safeguards against the deleterious consequences of the ribonucleotide excision by TOP1, including preventing the formation of TOP1-DNA adducts and associated cancer signature mutations. Moreover, our work also sheds light on the mechanism of PARP inhibitor cytotoxicity.

## Results

### Human TOP1 is prone to secondary cleavage and slippage mutations at genomic rNMP sites

TOP1 cleaves at a genomic rNMP site and generates a 2’, 3’-cyclic phosphate (CP) terminated nick.^8,23^ First shown in yeast Top1, a second cleavage by TOP1 a few nucleotides (2-5 bp) upstream from the CP nick may occur, which leads to the formation of TOP1cc and the release of the CP-containing segment (**Fig 1a**).^24^ When a duplex DNA substrate with a single embedded ribonucleotide was treated with human TOP1 in a 20-min end point analysis, interestingly, the predominant product type (as much as 60% of the total resulting species) was the TOP1cc product, with a lesser amount of the CP nick-product (**Fig 1b)**. To determine whether the TOP1cc was formed at the initial or the secondary cleavage site, the TOP1cc product was treated with proteinase K, followed by yeast tyrosyl-DNA phosphodiesterase 1 (Tdp1) and DNA 3’-phophatase (Tpp1), which together remove the TOP1 adduct and its bridging 3’-phosphate.^25,26^ These treatments generated a 25-nt product, corresponding to a secondary cleavage at 2-bp upstream of the CP nick (**Fig 1c**). In a time-course analysis, the amount of TOP1cc product increased over time, while the amount of CP nick product reduced slowly (**Supplementary Fig 2a**). We surmise that TOP1 is prone to secondary cleavage rather than ligation following the CP nick formation. Indeed, when TOP1 was titrated directly against a CP nick substrate, a clear preference for secondary cleavage was revealed, with TOP1cc representing up to 67% of the resulting species and a lesser amount of ligation product (up to 25% of the total products) (**Fig. 1d**). To further determine whether the ligation product was derived from direct CP nick ligation or slippage ligation, we tested human TOP1, with the yeast Top1 (yTop1) as a control, on a similar but shorter 45 bp CP nick substrate for a better separation of these two ligation products (**Fig 1e**). Interestingly, while a nearly equal amount of CP nick ligation and slippage ligation were observed for the human TOP1, yTop1 showcased a clear preference for CP nick ligation (over 3-fold more than the deletion product). Given the difference between human and yeast TOP1, we further compared their slippage mutation potential in the duplex substrate with an embedded ribonucleotide. In a time course analysis, human TOP1 again showcased an increased ability to generate deletion mutations compared to yTop1 (**Supplementary Fig 2b**). Notably, more substrate remained with yTop1 than with human TOP1 (∼69% versus 29% at 20 nM, respectively). This difference in remaining substrate is not the result of a reduced ability of yTop1 to cleave rNMP, since both proteins displayed comparable activities on a suicide fork substrate that does not allow for ligation once the initial cut is made (**Fig. 1f**). Thus, the drastic difference between human and yeast TOP1 activity on embedded rNMP is likely because yTop1 prefers ligation at the CP nick site, while human TOP1 prefers the secondary cleavage.

**Figure 1:**
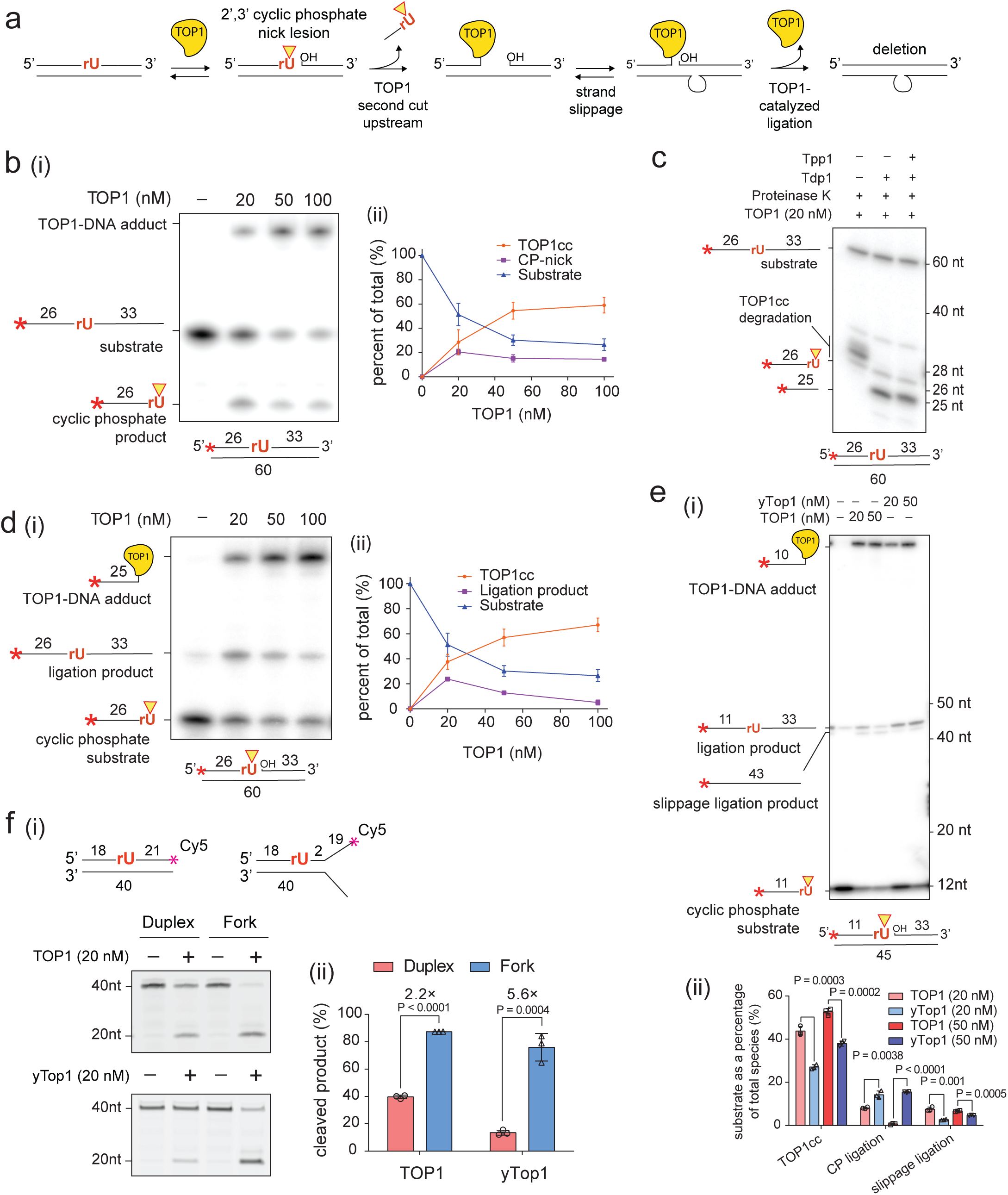
Human TOP1 is prone to secondary cleavage and TOP1cc formation at genomic rNMP sites. a) Diagram showcasing how TOP1 processes genomic ribonucleotides (rNMPs), and how this processing can lead to deletions in the context of short tandem repeats. b) (i) The titration of TOP1 against a 5’-^32^P-labeled duplex DNA containing a single embedded rNMP was analyzed in a denaturing gel, (ii) with the quantification of the resulting species. c) TOP1cc products were processed with Proteinase K, Tdp1, and Tpp1 to reveal the TOP1 cleavage site. d) (i) TOP1 was titrated against a 5’-^32^P-labeled CP nick substrate, (ii) with the quantification of the resulting species. e) (i) human TOP1 and yeast TOP1 (yTop1) were titrated against a CP nick substrate to examine their preference between CP nick ligation and slippage ligation. (ii) Quantification of the different products of TOP1 and yTop1. f) (i) 3’-Cy5-labeled duplex substrate and cleavage suicide substrate (fork) were tested with either human TOP1 or yTop1. (ii) The quantification of cleavage products.

### PARP1 regulates TOP1 ribonucleotide processing by suppressing its secondary cleavage

The secondary cleavage by TOP1 is directly linked to slippage mutations. The mutagenic nature of human TOP1 prompted us to probe its regulation for mutation suppression. Absent in yeasts, PARP1 is a first responder in single strand break repair in higher eukaryotes and has been reported to physically interact with TOP1, a known target for PARylation.^27–29^ More recently, TOP1 cleavage of genomic ribonucleotides was shown to generate PARP-trapping lesions,^22^ which motivated us to ask whether PARP1 is playing a role in regulating ribonucleotide excision by TOP1. PARP1 senses and is activated by DNA nicks. We first tested PARP1 in an electromobility shift assay (EMSA) using a mixture of equal amount of clean nick and CP nick substrates labeled with Cy5 and Cy3 respectively (**Supplementary Fig. 3a**). These results suggest that PARP1 recognizes both substrates equally well.

Interestingly, the addition of PARP1 to the TOP1 rNMP cleavage reaction drastically increased the amount of CP nick product (∼3-fold), while reducing the amount of TOP1cc (∼ 7-fold) (**Fig 2a**). Addition of NAD^+^, the cofactor necessary for PARP1-catalyzed PARylation, exacerbated this change, biasing nearly 98% of the products towards CP nick. Surprisingly, the presence of NAD^+^ in the reaction significantly increased the amount of remaining substrate (∼80%) likely due to the inhibition of the initial rNMP cleavage, and not the promotion of CP nick ligation, since PARP1 inhibits both TOP1-catalyzed secondary cleavage and ligation when testing the CP nick substrate (**Fig 2b**). Importantly, substitution of PARP1 with PARP1-E988K, a catalytic dead mutant of PARP1, showed a similar effect as PARP1 without NAD^+^ (**Supplementary Fig. 3b, c**), confirming PARP1 suppresses TOP1 secondary cleavage in both a catalytic-dependent and -independent manner, while it down-regulates TOP1-catalyzed initial rNMP cleavage only in a catalytic-dependent manner. To summarize, we surmise that upon the initial rNMP cleavage, PARP1 appears to protect the CP nick product from any further action by TOP1. The activated PARP1 also down-regulates TOP1 from further rNMP cleavage, but only in a catalytic-dependent manner.

**Figure 2:**
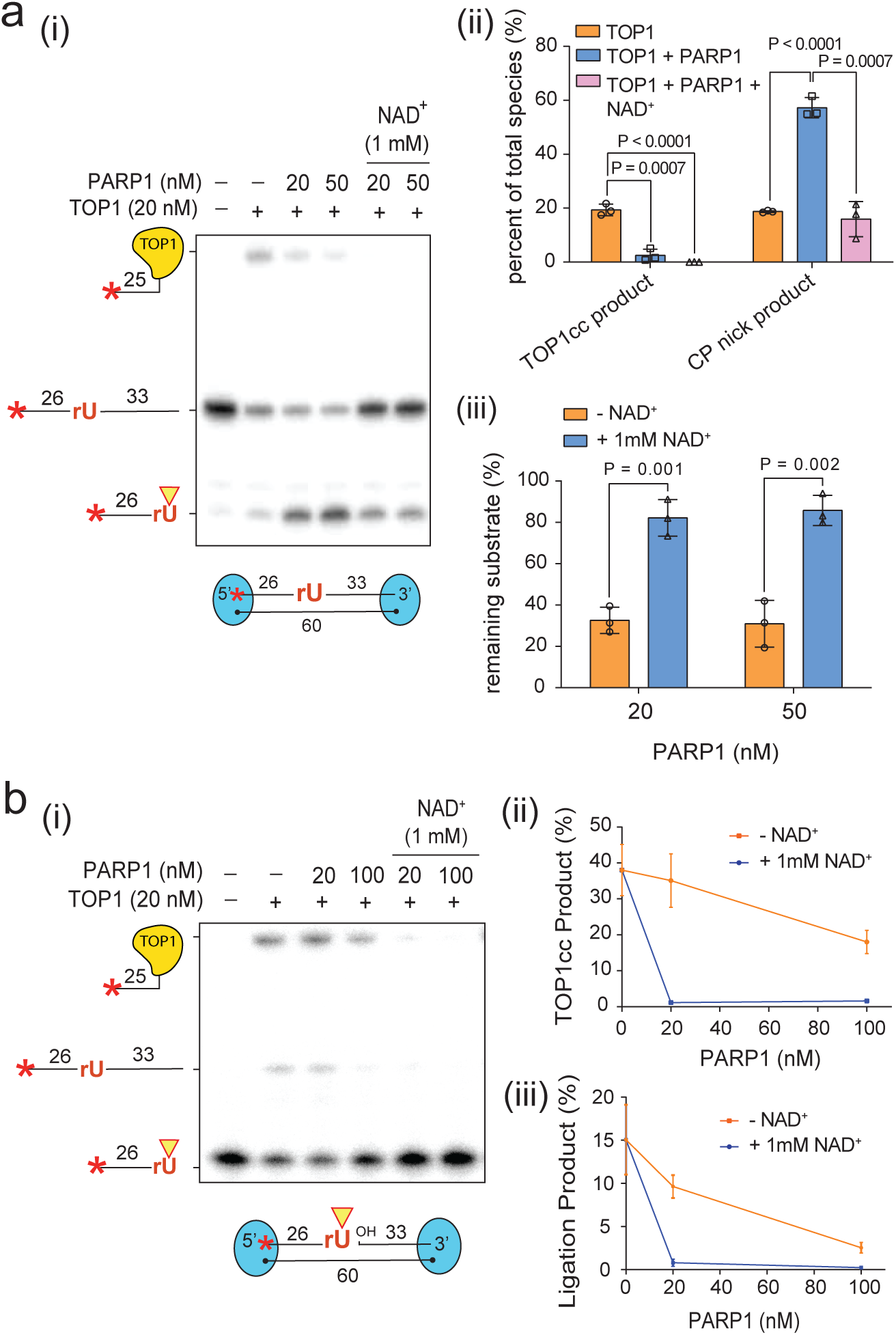
PARP1 suppresses the secondary cleavage by TOP1 at genomic ribonucleotides. a) (i) TOP1 ribonucleotide cleavage assay in the presence of PARP1 with or without NAD^+^. To prevent PARP1 from binding, both ends of the substrate were occluded by biotin-bound streptavidin as diagramed with blue ovals. (ii) The quantification of the CP nick and TOP1cc products at 50 nM PARP1 concentration and (iii) of the remaining substrate are shown. b) (i) The processing of the CP nick substrate by TOP1 was assessed in the absence or presence of PARP1 with or without NAD+, with both sides of the substrate occluded with streptavidin. (ii-iii) The quantification of the TOP1cc product and of the ligation products are shown.

### PARP1 knockout increases the slippage mutation rate due to increased TOP1 secondary cleavage

Our findings of PARP1 inhibition of TOP1 secondary cleavage strongly predict a role of PARP1 in suppressing slippage mutations. To test this hypothesis and ascertain TOP1 regulation by PARP1 in cells, we adopted a slippage mutation reporter HeLa cell line that a 2-bp slippage deletion will correct a frameshift mutation and restore puromycin resistance of the cells (kind gift from Dr. Andrew P. Jackson, **Fig 3a**). We knocked out PARP1 in both the parental reporter cell line and its derived RNASEH2A knockout cell lines (**Supplementary Fig 4a**). As expected from previous studies, RNASEH2A knockout significantly increased the slippage mutation rate (∼14-fold) compared to the wild-type cell line (Fig 3a-i). Consistent with our hypothesis, PARP1 knockout alone increased the slippage mutation rate by over 7-fold when compared to the wild-type. DNA sequencing of puromycin resistant clones from the PARP1 knockout cell line confirmed all 10 clones picked were 2-bp slippage events at STRs (**Fig 3a-ii, Supplementary Table 4**). Being previously reported as having G2/M arrest,^22^ RNASEH2A PARP1 double knockout cell line displayed a substantial slow growth phenotype, rendering the reporter assay technically challenging (**Fig 3b**). In addition to the homozygous RNASEH2A knockout, a partial deletion of RNASEH2A, namely RNASEH2A^+^, was also reported with around 50% loss of RNase H2 activity and causes a minor increase of slippage mutation rate.^12^ The knocking out of PARP1 in RNASEH2A^+^ led to an over 12-fold increase in the slippage mutation rate over the wild-type, above 5-fold over RNASEH2A^+^ and almost 2-fold over PARP1-KO (**Fig. 3a-i**). Hence, the impacts of RNASEH2A deficiency and PARP1-KO on slippage mutation rate are additive. Interestingly, the inhibition of PARP1 through PARP inhibitor talazoparib did not significantly change the mutation rate observed in the wild-type reporter cell line (**Supplementary Fig 4b**). This inhibition of deletions even in the absence of the catalytic activity of PARP1 indicates that PARP-trapping might compensate by physically impeding the TOP1 secondary cleavage.

**Figure 3:**
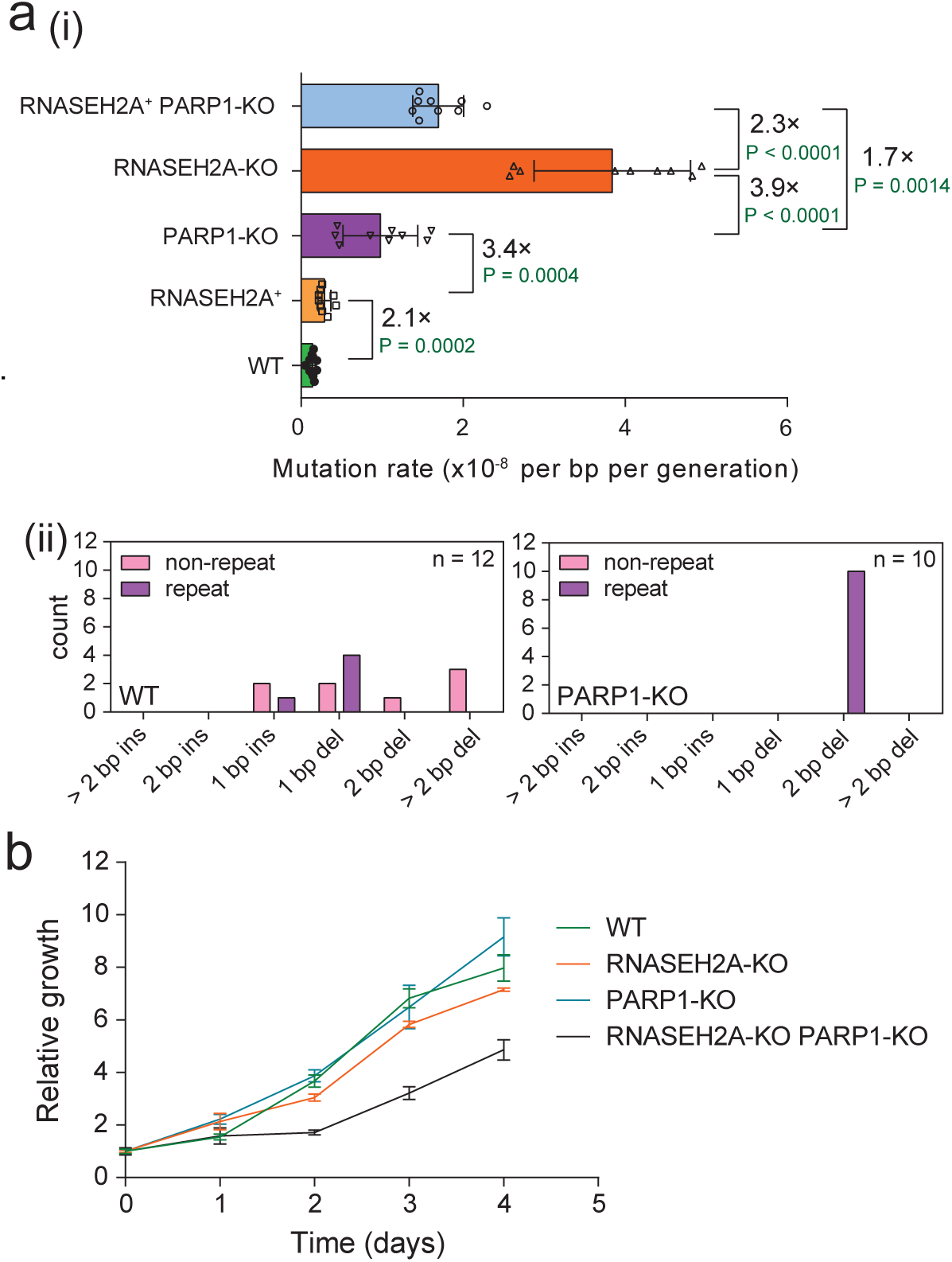
PARP1 prevents deletions at short tandem repeats, and its simultaneous deletion with RNASEH2A generates a slow growth phenotype. a) (i) Fluctuation assay reporting on the mutation rate of the different HeLa reporter strains as indicated. (ii) Quantification of the mutation types observed in sequenced clones of the WT and PARP1-KO strain showcases the prevalence of 2 bp deletions in the absence of PARP1. The n reflects the total number of indels identified in the total sequenced clones. b) Growth curve showcasing the delayed growth of the RNASEH2A-KO PARP1-KO cell line.

### Slow growth of RNASEH2A PARP1 double knockout cell line due to TOP1cc accumulation and its suppression by TDP1 overexpression

Based on our biochemically derived model, we suspected that the markedly reduced proliferation and G2/M accumulation in RNASEH2A-KO PARP1-KO cells is the result of excessive TOP1 secondary cleavage and thus the accumulation of stable TOP1cc. Navitoclax, a BCL2 inhibitor (BCL2i) that blocks the apoptosis suppressor BCL2, has been shown to enhance the cytotoxicity of the TOP1-trapping agent camptothecin in tumors, likely due to the pro-senescent role of BCL2.^30–34^ Consistent with our hypothesis of TOP1cc accumulation, the RNASEH2A PARP1 double knockout cells demonstrated hypersensitivity to navitoclax (**Fig. 4a**). To confirm the TOP1cc accumulation in the double knockout cells, we performed a RADAR (rapid approach to DNA adduct recovery) assay designed to detect the amount of TOP1cc.^35,36^ Our results showed that the double knockout cell line has a 3-fold increase in the formation of TOP1cc when compared to wild-type and significantly higher than the single knockouts of either RNASEH2A or PARP1, with a difference of ∼1.5 and 2-fold respectively (**Fig 4b**). To test if the buildup of TOP1cc was the cause for the slow growth, we overexpressed TDP1, a protein that is crucial for TOP1cc removal in cells,^25^ which almost completely rescued the growth defect of our double knockout cells (**Fig 4c**). Moreover, the treatment of the double knockout cell lines with a TDP1 inhibitor (TDP1i), CD00509,^37^ led to an over 60% reduction in viability (**Fig 4c**). This TDP1i sensitivity was further aggravated by BCL2 inhibition, affirming TOP1cc as the toxic lesion slowing the proliferation of the RNASEH2A-KO PARP1-KO cells (**Fig. 4d**). These results confirmed the role of PARP1 in the suppression of the detrimental TOP1 secondary cleavage at genomic rNMP sites in cells.

**Figure 4:**
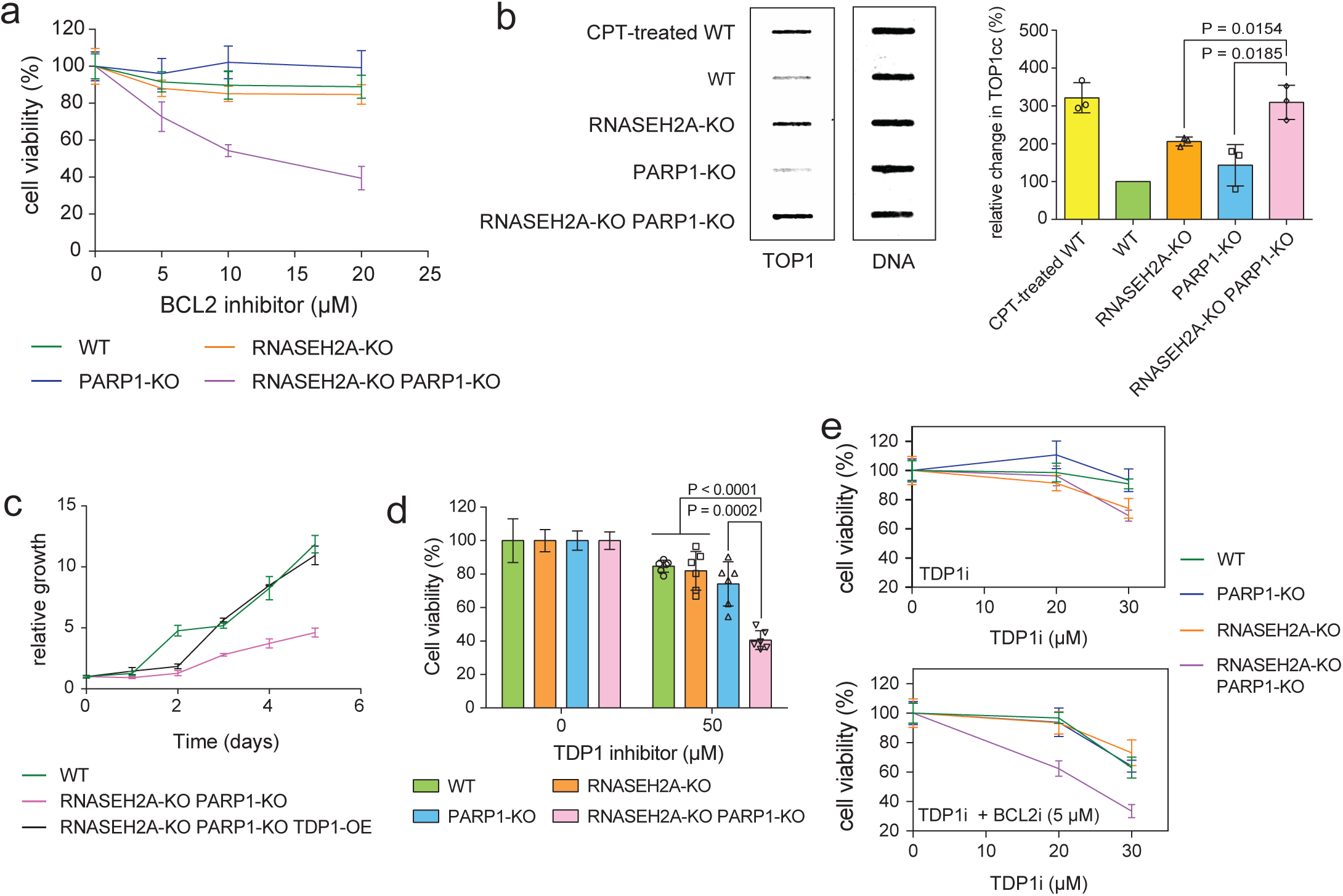
Concurrent deletion of RNASEH2A and PARP1 generates a detrimental accumulation of TOP1cc. a) Sensitivity of WT, RNASEH2A-KO, PARP1-KO and RNASEH2A-KO PARP1-KO cell lines to the BCL2 inhibitor (BCL2i) navitoclax was determined by the MTT viability assay. b) Slot blot of the RADAR assay showcasing the elevated accumulation of TOP1cc in the RNASEH2A-KO PARP1-KO cell line. Samples from treating WT cells with 10 μM of the TOP1-trapping agent camptothecin (CPT) were included as a positive control. The DNA loading control was stained with Sybr Gold. c) Growth curve showing that the overexpression of TDP1 (TDP1-OE) rescues the delayed growth in the RNASEH2A-KO PARP1-KO cell line. d) Viability measured by MTT assay after TDP1 inhibitor, CD00509, (TDPi) treatment demonstrates hypersensitivity of the RNASEH2A-KO PARP1-KO cell line to TDP1 inhibitor. e) MTT viability assay showcasing the synthetic effect of TDP1i and BCL2i treatments to the viability of the RNASEH2A-KO PARP1-KO cell line.

### A feedback loop of CP nick formation, PARP1 activation and TOP1 inhibition

In the reconstituted system, catalytically, PARP1 inhibits TOP1-catalyzed secondary cleavage and CP nick ligation, as well as the initial rNMP cleavage. Analysis of the PARylation status of the proteins in the reaction containing the embedded ribonucleotide substrates confirmed that TOP1 was PARylated in the presence of NAD^+^ (**Fig 5a**, **5b**), which is consistent with previous proteomic studies.^28,29^ Notably, the presence of TOP1 stimulated PARP1 auto-PARylation in the presence of rNMP embedded duplex DNA. Treating the reactions with Poly (ADP-ribose) glycohydrolase (PARG), an enzyme that removes the ribose chains from proteins and leaves a single ADP-ribose in the modified residues,^38^ further confirmed TOP1 PARylation and revealed an over 5-fold increase in PARP1 auto-PARylation when TOP1 was present (**Fig 5b**). This stimulation in auto-PARylation is likely the result of the formation of CP nick that then activates PARP1, as this stimulation was not observed with the catalytically inactive TOP1-Y723F (**Fig 5c**).^39,40^ Therefore, the CP nick formed after initial rNMP cleavage by TOP1 activates PARP1, which then PARylates TOP1 and inhibits TOP1-catalyzed cleavage of additional rNMP sites (diagram shown in **Fig 5d**).

**Figure 5:**
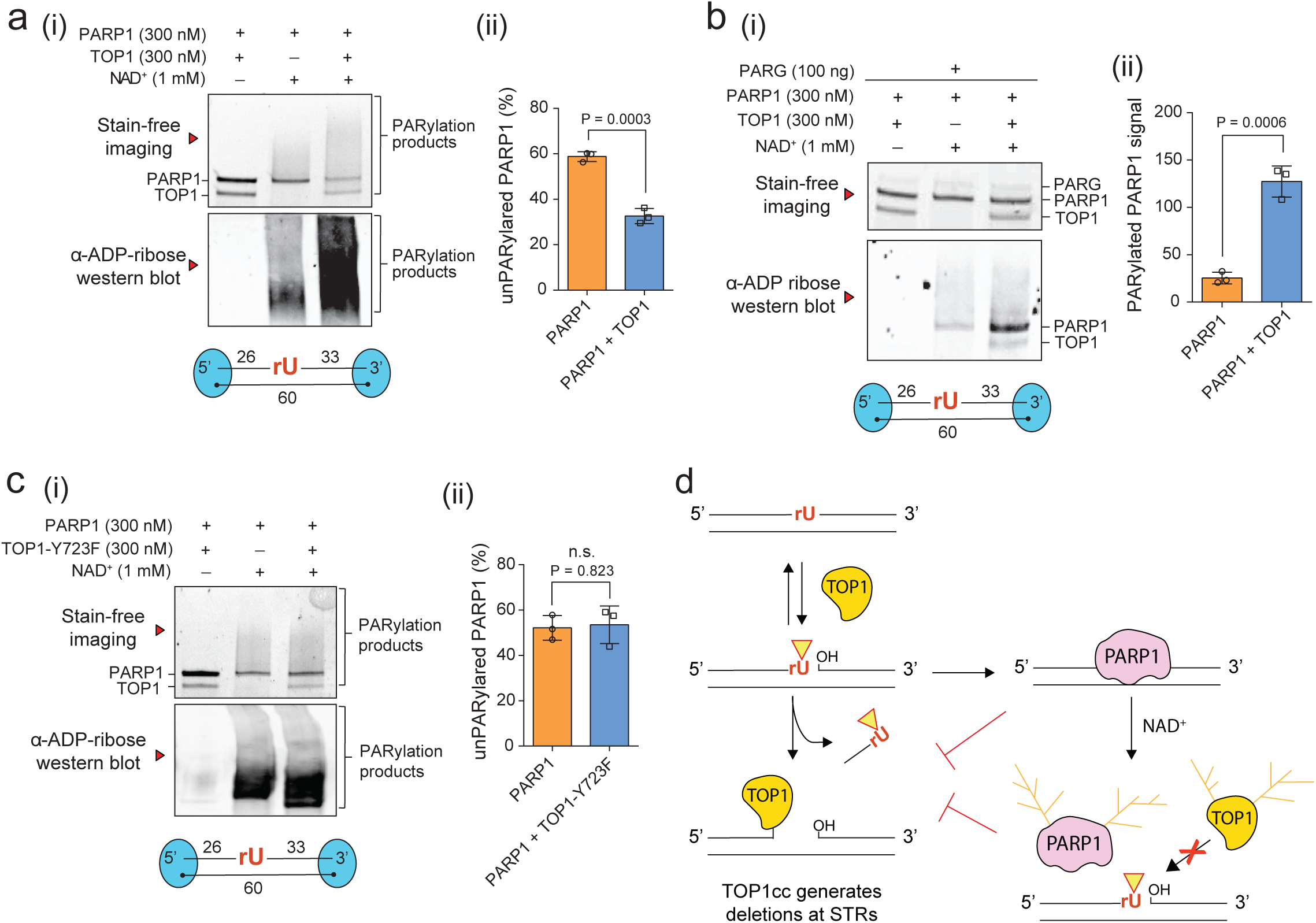
TOP1 initial cleavage at genomic ribonucleotides stimulates PARylation by PARP1. a) (i) PARP1-catalyzed auto-PARylation and PARylation of TOP1 in the presence of the duplex substrate with a single embedded ribonucleotide *in vitro*. Both DNA ends were occluded by biotin-bound streptavidin. (ii) Quantification of the amount of unPARylated PARP1 in the absence or presence of TOP1. b) (i) The *in vitro* PARylation assay using the ribonucleotide-containing substrate where the products were further treated with PARG, leaving single ADP-ribosyl modification that allows for precise determination of protein PARylation levels (ii) Quantification of PARylation levels of PARP1 in the absence or presence of TOP1. c) (i) *in vitro* PARylation assay with the ribonucleotide-containing substrate using the catalytic dead mutant TOP1-Y723F. Quantification of the amount of unPARylated PARP1 in the absence or presence of TOP1-Y723F. d) Diagram showcasing model on how PARP1 prevents the secondary cleavage by TOP1 at ribonucleotide sites both in a catalytically-independent (by physically restricting TOP1) and - dependent manner (through the inhibitory PARylation of both TOP1 and itself).

### Both DNA binding and TOP1 interaction are essential for PARP1 regulation of TOP1 ribonucleotide processing

In addition to catalytical regulation of TOP1 by PARP1, we also found that PARP1 inhibits CP nick cleavage and ligation by TOP1 in a non-catalytic manner. However, the DNA binding domain (DBD) harboring zinc finger motif 1, 2 and 3 was unable to suppress the secondary cleavage by TOP1, nor does the N-terminal truncation that removes the DNA binding domain (DBD) of PARP1 (**Fig 6a, b**). Hence, the binding of PARP1 to CP nick is necessary, but not sufficient, for TOP1 secondary cleavage inhibition. This led us to ask if the reported physical interaction between PARP1 and TOP1 is also essential for TOP1 regulation.^27^ To map the interaction between PARP1 and TOP1, we generated a series of PARP1 truncations, starting from the C-terminus, and tested them for TOP1 interaction using *in vitro* pull-down assays (**Fig 6c**). Interestingly, the removal of its adjoining helical domain (HD), an autoinhibitory motif that locks PARP1 activity before binding to its substrate DNA, completely disrupted PARP1 and TOP1 interaction (**Fig. 6d**).^41,42^ These results suggest the HD domain as a key interface between PARP1 and TOP1.

**Figure 6:**
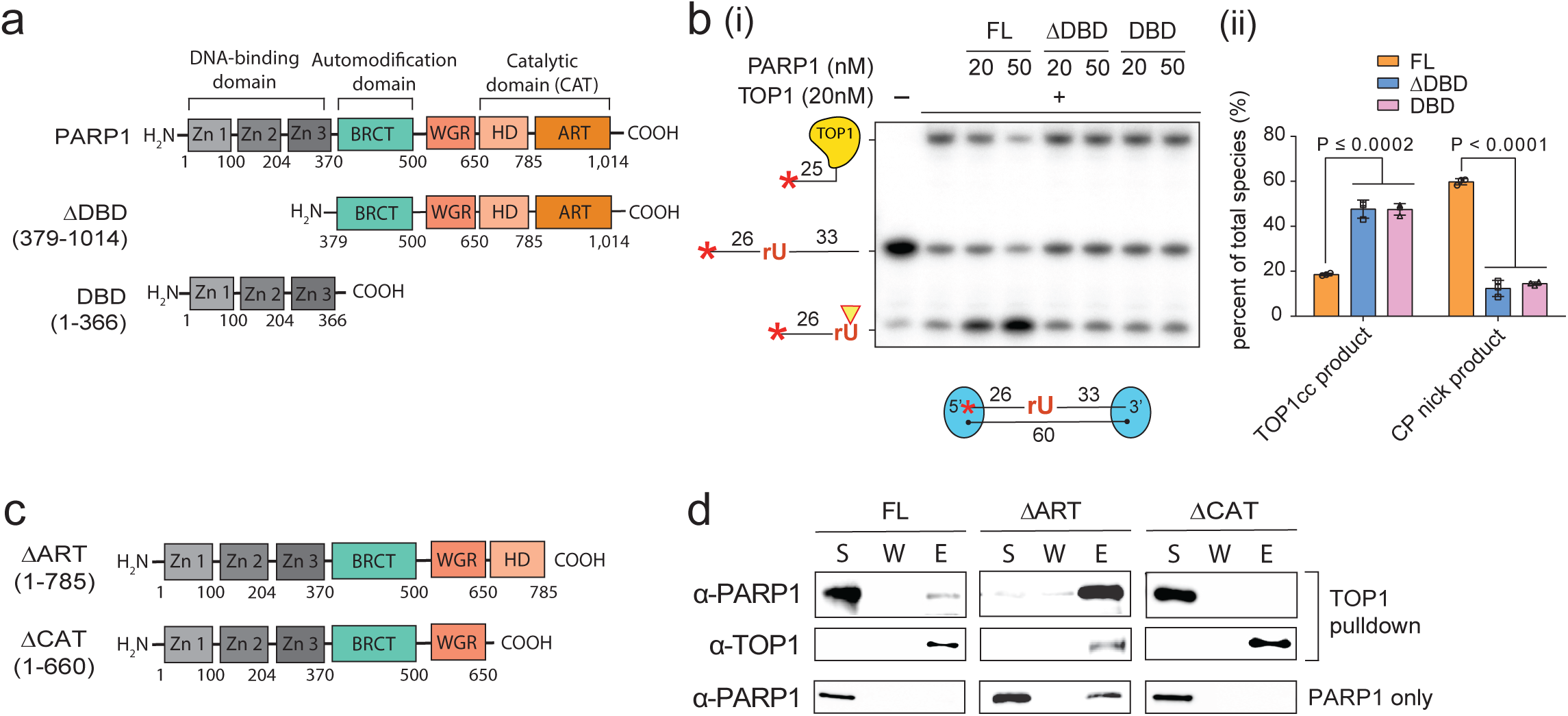
The DNA-binding activity of PARP1 and its physical interaction with TOP1 are important for the prevention of the TOP1 secondary cleavage. a) Diagram showing the different domains of the full-length (FL) PARP1 and the two PARP1 truncations (PARP1-ΔDBD and PARP1-DBD) tested in the TOP1 ribonucleotide cleavage assay (i) The TOP1 ribonucleotide cleavage assay in the absence or presence of the full-length PARP1-FL, PARP1-ΔDBD or PARP1-DBD. (ii) Quantification of the cyclic phosphate (CP) product and the TOP1cc product for the reactions with PARP1-FL, PARP1-ΔDBD and PARP1-DBD at 50 nM concentration. c) Domain diagram of N-terminal truncations of PARP1 (PARP1-ΔART and PARP1-ΔCAT) tested for TOP1-binding. d) The *in vitro* pulldown of FLAG-tagged TOP1 against the full-length PARP1 and PARP1 N-terminal truncations (ΔART and ΔCAT). Reactions without TOP1 were included as controls to assess non-specific binding of PARP1 to the anti-FLAG M2 resin.

### PARP1-R735E mutation partially disrupts its interaction with TOP1 and fails to fully restore PARP inhibitor sensitivity in PARP1-KO cells

It has been reported that portions of the core domain and C-terminus region TOP1 participate in PARP1 interaction.^27^ Using Alphafold 3, we modeled the complex between TOP1-ΔN200, lacking the N-terminal disordered region of TOP1, and the PARP1 HD domain. Notably, among the top four out of the five models generated, the TOP1-HD interface consistently involved the same surface of the HD domain, which contains a cluster of conserved basic residues (K695, K700, R704, and R735) (**Supplementary Fig 5a, b**). Inspired by this finding, we generated three mutants that had their positive residue substituted with the negative charged glutamic acid to assess their binding to TOP1. Our pull-down results showed that both the R735E and the concurrent K700E/R705E mutant largely abolished binding with TOP1 (**Fig 7a**). We chose to focus on the PARP1-R735E mutant, as this purified mutant is comparable to the wild-type PARP1 in nick binding and auto-PARylation, and sensitive to PARP inhibitor (PARPi), talazoparib, *in vitro* (**Supplementary Fig 5c and d**). Moreover, when expressed in PARP1-KO U2OS cell lines, PARP1-R735E retained its ability to rescue the PARP1-KO from camptothecin (CPT) sensitivity,^43,44^ while displaying reduced interaction with TOP1 (**Supplementary Fig 5e, f and Fig 7b**). This deficiency in interaction of the R735E mutant with TOP1 appears to affect the inhibition of TOP1 secondary cleavage as TOP1cc formation was reduced nearly 3-fold with the R735E mutant compared to the wild-type PARP1 (at 50 nM PARP1 concentration) (**Fig 7c**). The R735E mutant also showcased a significant decrease in PARylation-dependent inhibition of TOP1 secondary cleavage and initial rNMP cleavage (**Fig 7d**). Importantly, the R735E mutant desensitized cells to PARP inhibitor talazoparib when compared to the cells expressing the wild-type PARP1, with a difference in half-maximal effective concentration (EC_50_) of almost 4-fold (**Fig 7e**). This result suggests that the interaction with TOP1 might be crucial for the engagement of PARP1 at sites of ribonucleotide cleavage by TOP1. Overall, our findings highlight the relevance of the physical interaction between TOP1 and PARP1 for the regulation of ribonucleotide processing and PARP trapping.

**Figure 7:**
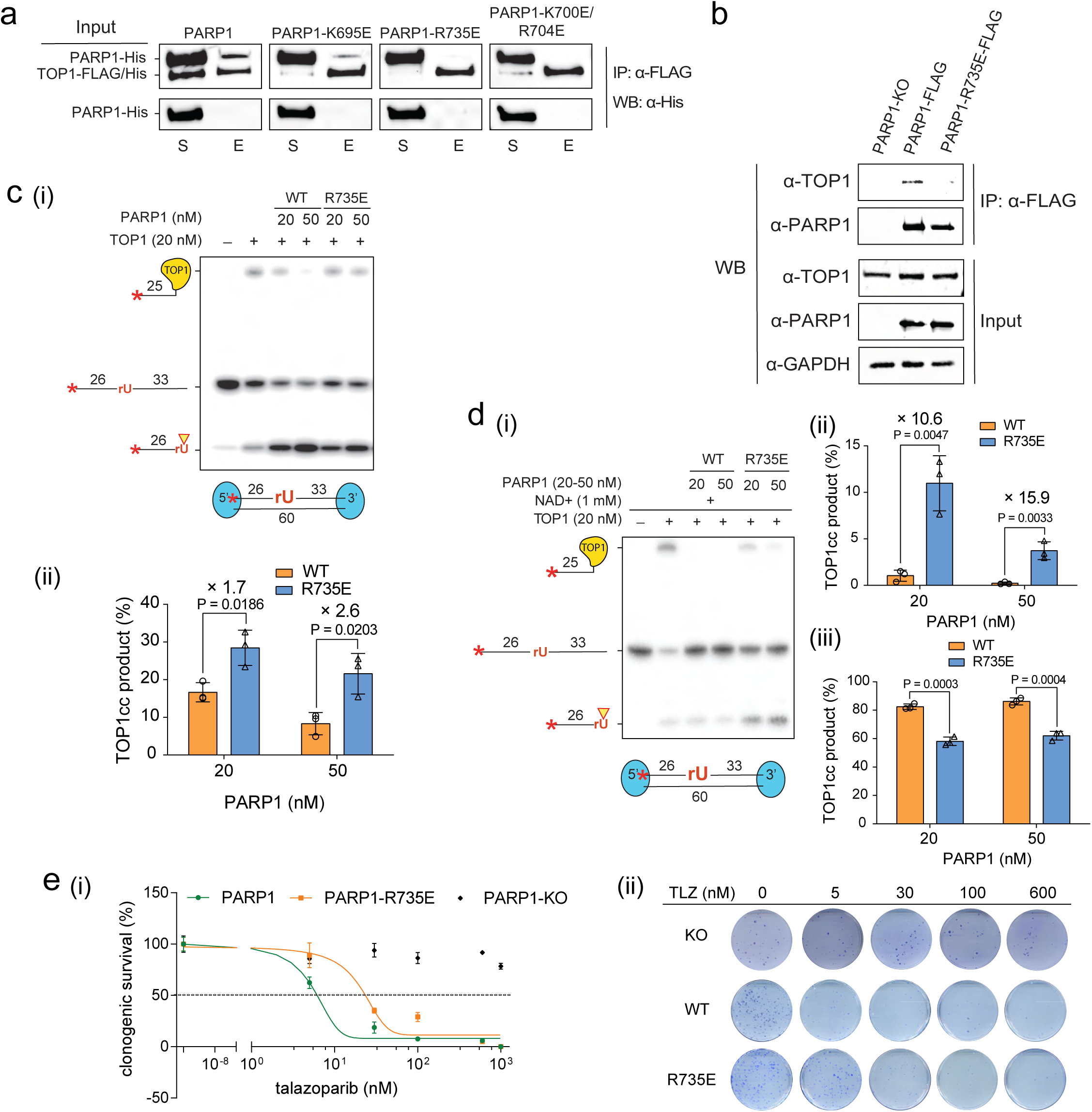
Characterization of PARP1-R735E mutant in TOP1 interaction, the regulation of TOP1 ribonucleotide cleavage and sensitivity to PARP inhibitor, talazoparib. a) The *in vitro* anti-FLAG-pull-down assay to test interaction of TOP1 with PARP1 mutants. The membrane was blotted with anti-His antibody for the detection of TOP1 and PARP1, both His-tagged. The supernatant (S) and elution (E) of the pull-down assay are shown. b) co-immunoprecipitation analysis of interaction between TOP1 and PARP1 in cells, where anti-FLAG M2 resin was used to immunoprecipitate either FLAG-tagged PARP1 or PARP1-R735E mutant stably expressed in U2OS PARP1-KO cell line to test their interaction with TOP1. c) (i) The TOP1 ribonucleotide cleavage assay in the absence or presence of PARP1 or PARP1-R735E. (ii) Quantification of the TOP1cc product from the reaction with either WT or the R735E mutant PARP1. d) (i) TOP1 ribonuclease assay in the presence of NAD^+^ comparing TOP1 inhibition by the WT PARP1 and the R735E mutant. e) (i) Clonogenic survival curve testing the sensitivity to talazoparib (TLZ) treatment of PARP1-KO U2OS cell lines expressing either no PARP1, WT PARP1, or PARP1-R735E (normalized to untreated cells and fitted using a nonlinear least-squares fit to a three-parameter dose–response model). (ii) A representative plate for each strain at the different TLZ concentrations as indicated.

## Discussion

The processing of genomic ribonucleotides by TOP1 results in three major products that can compromise genome integrity: the CP nick, the TOP1cc, and the deletions at STRs. The human genome is densely populated with tandem repeats, which together account for approximately 3% of the genome, with dinucleotide repeats alone accounting for approximately 0.5% of the total genomic sequence.^45^ This abundance of STRs, as well as the mutagenic nature of human TOP1, requires a mechanism to safeguard the genome in cells. Our findings reveal that PARP1 plays a crucial role in preventing these deletions by inhibiting TOP1 secondary cleavage at sites of genomic rNMPs. It does this through both catalytic-independent and -dependent mechanisms. Non-catalytically, PARP1 physically impedes the TOP1 secondary cleavage, which requires both nick binding and TOP1 interaction. Catalytically, following its activation upon binding to CP nick, PARP1 is capable of PARylating TOP1, which likely inhibits several activities of TOP1, including the rNMP cleavage, the secondary CP nick cleavage and CP nick ligation.

The concurrent deletion of both PARP1 and RNASEH2A led to a pronounced accumulation of TOP1cc. This is likely because failure of genomic rNMP removal by ribonucleotide excision repair shifts processing to TOP1-mediated cleavage, while the absence of PARP1 increases susceptibility of TOP1 to secondary cleavage. The resulting accumulation of trapped TOP1 proved to be deleterious to the growth of PARP1 and RNASHE2A double knockout cells, as it can be rescued by TDP1 overexpression. Notably, these cells were hypersensitive to BCL2 inhibitors, which induce apoptosis in senescent cells by blocking BCL2-mediated suppression of programmed cell death.^32,33^ Consistent with our findings, TOP1cc accumulation has been linked to cellular senescence through the cGAS-mediated inflammatory senescence pathway.^31,34^

An important aspect of genomic rNMP removal by TOP1 is its link to inhibitor-induced PARP trapping.^22^ *In vitro*, purified PARP1 binds CP nicks with an affinity comparable to that for clean nicks, supporting CP nicks as a relevant PARP1 trapping lesion. Notably, whereas PARP1 binding to clean nicks has little effect on ligation by ligase III,^46,47^ binding to CP nicks blocked both TOP1-mediated CP nick ligation and CP nick cleavage. Thus, PARP1 trapping at CP nicks may impede their repair and cause replication fork collapse. Importantly, treatment with a PARP inhibitor had little effect on mutation rates, suggesting that trapped PARP1 efficiently blocks TOP1 secondary cleavage in a catalysis-independent manner. In contrast, PARP1-R735E, which retains DNA binding and catalytic activity but is defective in TOP1 interaction and suppression, partially rescued PARP inhibitor-induced cytotoxicity. This means that the physical interaction with TOP1 is relevant for engaging PARP1 at sites of ribonucleotide cleavage, such as due to recruitment of PARP1 by TOP1 or a handoff of DNA at the cleavage site between TOP1 and PARP1. Together, these findings establish a potential mechanism underlying PARP inhibitor cytotoxicity and its connection to TOP1 regulation during genomic rNMP removal.

Aicardi-Goutières syndrome (AGS), a neurodevelopmental disorder, and systemic lupus erythematosus (SLE) are both RNase H2-associated autoimmune disorders characterized by hyperactivation of the innate immune response in cells.^48^ In both cases, this immune response is thought to arise from the genomic instability that occurs in the presence of a dysfunctional RNase H2 complex.^6,48–50^ An improved understanding of genomic rNMP processing in the absence of RNase H2 in human cells will provide a framework to identify the source of the genomic instability that leads to this innate immune response in AGS and SLE. Given our findings that PARP1 limits the accumulation of TOP1cc in the absence of functional RNase H2, it would be interesting to assess which of these rNMP-associated lesions (CP nick and TOP1cc), if any, participate in the emergence of innate immune response in these disorders. TOP1cc, for example, has already been linked to cGAS activation.^31^ Although most cells in the nervous system are post-mitotic, proliferative immune cells such as microglia can drive neuronopathic immune responses even when they are the only cells harboring AGS genetic mutations.^51^ Furthermore, mice models of AGS mutations showcased increased phagocytosis activity in microglia, as well as elevated levels of type I interferon mainly within microglia.^52^ These observations are particularly relevant because replicating cells are expected to freshly incorporate ribonucleotide and may therefore be more susceptible to genome instability arising from TOP1-mediated rNMP processing.

## Methods

### Plasmids and oligonucleotides list

The description of all the plasmids utilized has been provided in the **Supplementary Table 1**. The oligonucleotides used to generate the substrates for biochemical assays are provided in the **Supplementary Table 2**.

### Expression and purification TOP1, yTop1 and TOP1-Y723F from insect cells

The coding sequence for TOP1 was amplified from TOP1 cDNA (Dharmacon Reagents) and cloned into a modified pFASTBac-HTB vector and fused in frame to the FLAG and His_6_ tags within the vector.^53^ The construct was transformed into DH10Bac *E. coli* strain (Invitrogen) to generate the bacmid, which was used to infect SF9 insect cells for TOP1 expression. To purify TOP1, the insect cell pellet (∼15g from 1 L culture) was resuspended in 50 mL K500 buffer (500 mM KCl, 20 mM KH_2_PO_4_, pH 7.4, 10% glycerol, 0.5 mM EDTA, 0.01% NP-40, 1mM β-mercaptoethanol) with a cocktail of protease inhibitors (aprotinin, chymostatin, leupeptin, and pepstatin A at 5 μg/ml each, and 1 mM phenyl-methylsulfonyl fluoride). The resuspended cells were disrupted by sonication for 1 minute, and the lysate was clarified by centrifugation (20,000 g for 20 minutes). The supernatant was then loaded onto a 3 mL Ni-NTA column (Thermo Fisher) using a BioRad Econo pump. The column was washed with 50 mL K500 buffer with 10 mM imidazole. TOP1 was eluted using 20 mL K500 buffer with 250 mM imidazole. The eluate was then incubated with 0.5-2 mL anti-FLAG M2 agarose resin (Sigma) overnight. The following day, the beads were spun down (1000 g for 5 minutes), and the supernatant was removed. The matrix was washed with K500 buffer with an adjusted 0.1% NP-40. TOP1 was finally eluted by incubating 1.2 mL of K500 containing 200 μg/mL FLAG peptide for 1 hour. The eluate was filter-dialyzed to remove excess FLAG peptide. The purified Top1 protein (∼0.5 mg) was frozen in liquid nitrogen and stored at −80°C as small aliquots. To express TOP1-Y723F, the mutation was introduced via site-directed mutagenesis, and the protein was expressed similarly as WT, with a yield ∼3 mg. The coding sequence of yTop1 was amplified from yeast genomic DNA and cloned into the modified pFASTBac-HTB with a Flag tag and a His_6_ tag engineered at the amino-terminus. The same procedure was followed for the purification of yTop1 as reported.^54^

### Expression and purification of recombinant PARP1 and its mutants from *E. coli*

The pET28 plasmid expressing full-length PARP1 (Addgene #192635, a kind gift from John Pascal)^55^ was transformed into Rosetta 2 pLysS *E. coli* strain (Novagen) for protein expression. The cells were grown in 8 L of LB broth supplemented with 10 mM benzamide, and protein expression was induced at OD_600_=0.6 with 0.5 mM IPTG and 100 μM ZnSO_4_ was supplemented at the same time. After IPTG addition, protein expression was allowed to occur overnight at 18 °C. To purify PARP1, the bacterial pellet (∼15g) was resuspended in 50 mL N500 buffer (500 mM NaCl, 20 mM Tris, pH 7.0, 10% glycerol, 0.5 mM EDTA, 0.01% NP-40, 1mM β-mercaptoethanol) with a cocktail of protease inhibitors (aprotinin, chymostatin, leupeptin, and pepstatin A at 5 μg/ml each, and 1 mM phenyl-methylsulfonyl fluoride). The resuspended cells were disrupted by sonication for 10 minutes, and the lysate was clarified by centrifugation (20,000 g for 20 minutes). The supernatant was then split onto two gravity columns with 2 mL HisPur Ni-NTA resin each (Thermo Fisher). Subsequently, the column was washed with 50 mL K500 buffer with 20 mM imidazole. PARP1 was eluted using 20 mL N500 buffer with 250 mM imidazole. The eluate was then diluted to 150mM salt concentration and loaded into two gravity columns with 2 mL Heparin Sepharose 6 Fast Flow resin (Cytiva). Each column was then washed with 25 mL N150 (150 mM NaCl, 20 mM Tris, pH 7.0, 10% glycerol, 0.5 mM EDTA, 0.01% NP-40, 1mM β-mercaptoethanol). The protein in each column was then eluted with 10 mL N500, and the combined protein elution was concentrated in an Ultracel-30K concentrator (Amicon). The eluted protein was finally loaded into a Superdex 200 increase gel filtration column (Cytiva) connected to an AKTA pure chromatography system (Cytiva) and eluted with K500 buffer. The eluted PARP1 (∼1 mg) is free of DNA contamination and thus show minimal PARylation activity in the absence of DNA substrate. To generate PARP1 mutants, mutations were introduced via site-directed mutagenesis, and the proteins were expressed similarly as WT. The C-terminal truncations of PARP1 (ΔC785 and ΔC660) were purified in the same way as WT. For these truncations, we used two pET28 constructs from the Conaway lab (Addgene #174804 and # 173942).^56^ The ΔC785 truncation, specifically, requires 12 L of cells rather than 8 L. The construct for the PARP1 DBD-only truncation (N-366) was made by amplifying and cloning the DBD region of PARP1 into the pETDuet-1 plasmid. This truncation was purified in a similar fashion to the previous PARP1 proteins. The N-terminal truncation of the DBD (ΔDBD) was also cloned in a pETDuet-1 with His_6_ and FLAG tags and purified with Ni and anti-FLAG affinity chromatography (similarly as described with TOP1).

### Generating 2’, 3’-cyclic phosphate terminated (CP) nick substrates

To generate the 2’, 3’-cyclic phosphate terminated nicked duplex, the H4 oligonucleotide containing two consecutive ribonucleotides was treated with RNase I_f_ (New England Biolabs) for 5 minutes at 37°C and subsequently heat inactivated. This reaction yields ∼ 40-60% of 2’, 3’-cyclic phosphate terminated ends that are resistant to alkaline phosphatase (New England Biolabs). The remaining RNase I_f_ product corresponds to the 3’-phosphate-terminated oligonucleotide. To obtain a pure cyclic phosphate product, the reactions were treated with calf intestinal alkaline phosphatase and thermolabile exonuclease I (New England Biolabs) for 20 mins to convert the 3’-phosphate ends into hydroxyl terminated oligonucleotides and subsequently digest them. The reactions were then heat inactivated for 3 mins at 80°C, and the proteins were removed with oligo cleaning columns (Zymogen). The purity of our cyclic phosphate terminated product was assessed with alkaline phosphatase and T4 PNK treatment (New England Biolabs), in which the first enzyme does not process the cyclic phosphate while the latter can remove it.^57^ The product was then 5’-P^32^ labelled using minus 3-phosphatase T4-PNK (New England Biolabs) and cleaned using a Spin6 column (BIO-RAD). To make the final cyclic phosphate terminated nicked duplex, the product was annealed with H3 and H5 oligonucleotides via slow cooling from 75 °C to avoid the spontaneous hydrolysis of the cyclic phosphates. To make the shorter 45 bp CP nick substrate, the treated DNA was H6 and it was annealed to H7 and H5 oligonucleotides. Streptavidin in our reaction buffers will bind to the two biotinylated ends of the H3 or H7 oligonucleotide to prevent the engagement of PARP1 at the blunt ends of the substrate. To make the CP nick containing dumbbell substrate, the H12 oligonucleotide was treated with RNase I_f_ as described in this section and then annealed by immediate cooling on ice following heat denaturing at 95°C for 5 minutes to minimize inter-molecular annealing. The same annealing procedure was followed for the regular nick dumbbell substrate.

### TOP1 ribonuclease assay

For TOP1 ribonuclease assays, the indicated amount of TOP1 in the reaction buffer (20 mM Tris-HCl, pH 8.0, 2 mM MgCl_2_, 100 μM TCEP, 100 μg/mL BSA, 150 mM KCl, 20 μg/mL streptavidin) was incubated with 10 nM DNA substrate for 20 min at 37°C. The reaction was stopped by treatment with SDS (0.2% final). When determining the identity of the TOP1 adduct, the reaction was treated with proteinase K (0.5 mg/ml) before column purification (oligo clean & concentrator, Cat. No: D4060, Zymo Research), followed by treatment with both yeast tyrosyl-DNA phosphodiesterase 1 (Tdp1) and DNA 3’-phophatase (Tpp1). Equal volume of loading dye containing 85% formamide with 25 mM EDTA, and 1 μM unlabeled H1 DNA strand (same sequence as rNMP-containing oligonucleotide) was then added to reaction mixtures to prevent re-annealing of the radiolabeled strand. The mixture was heated at 75 °C for 5 min and cooled on ice for another 5 mins. The reactions were run in 12% denaturing polyacrylamide gel in TBE (Tris-Boric Acid-EDTA) buffer, followed by phosphor imaging analysis.

### PARylation assay

For *in vitro* PARylation assay, we followed a similar procedure as previously published, with slight modifications for the experiments that include the rU-containing substrate.^58^ Specifically, the auto-PARylation assays with the dumbbell substrates (1µM) were incubated for 10 mins in a buffer consisting of 20 mM Tris-HCl, pH 8.0, 2 mM MgCl_2_, 100 μM TCEP, and 150 mM KCl. For the PARylation assay using the rNMP containing substrate (10 nM), the reaction buffer was supplemented with 20 μg/mL streptavidin to ensure the occlusion of the two blunt ends of our substrate and thus preventing unwanted activation of PARP1 by double strand DNA ends. Other than the negative controls and NAD^+^ titrations, all the other reactions contain 1 mM NAD^+^. Before the addition of TOP1 or PARP1, the reactions were pre-incubated for 10 mins at 37°C to ensure streptavidin is bound to the substrate ends. Once the proteins were added, the reactions were incubated for 10 mins at 25°C. The reactions were stopped with 3x Laemmli SDS buffer and heated for 5 minutes at 95°C. The reactions were run in a Mini Protean TGX Stain-Free 4-15% SDS gel (BIO-RAD). The gels were first scanned with stain-free imaging in a BIO-RAD Gel Doc EZ imager and subsequently transferred onto a nitrocellulose membrane for western blot analysis with the Anti-Poly/Mono ADP-ribose rabbit monoclonal antibody (Cell Signaling, D9P7Z, 1:1,000 dilution). For detection, we used IRDye 800CW goat anti-rabbit antibody (LICOR, 926-32211, 1:10,000 dilution).

### DNA binding assay and binding competition assay

Dumbbell substrates used in the DNA binding assays were generated as described in the “Generating 2’, 3’-cyclic phosphate terminated substrates” subsection of the Methods. Either the CP nick dumbbell substrate, labeled internally with a Cy3 dye, or the clean nick dumbbell substrate, labeled internally with a Cy5 dye (or both substrates in the case of the competition assay) was incubated with indicated concentrations of PARP1 for 20 mins in a buffer containing 20 mM Tris-HCl, pH 8.0, 2 mM MgCl_2_, 100 μM TCEP, 100 μg/ml BSA, and 150 mM KCl. The reactions were resolved using a 6% native gel and imaged using a ChemiDoc MP imaging system (BIO-RAD).

### Cell Lines

All the human cell lines used are depicted in the **Supplementary Table 3**.

### *In vitro* affinity pulldown assay

To test for TOP1-PARP1 interaction, (His)_6_-tagged PARP1 (350 ng) was mixed with FLAG-(His)_6_-tagged TOP1 (750 ng) in 25 μL buffer (20 mM Tris-HCl, pH 8.0, 2 mM MgCl_2_, 100 μM TCEP, 100 μg/ml BSA, 150 mM KCl). The mixture was incubated with 25 μL anti-FLAG M2 agarose resin (Sigma) for 1 hour shaking at 25°C. Following incubation, the resin was washed twice with 400 μl of wash buffer (150 mM KCl, 20 mM KH_2_PO_4_, pH 7.4, 10% glycerol, 0.5 mM EDTA, 0.1% NP-40, 1mM β-mercaptoethanol), and 20 μl of 2% SDS was used to the elute proteins. The supernatant (S) and SDS eluate (E) fractions, 10 μl each, were resolved by 8% SDS-PAGE and immunoblotted with either HRP-conjugated anti-His antibodies or with their respective protein specific antibodies.

### Fluctuation assay

The engineered HeLa reporter cell lines including the WT, RNase H2A+, and RNase H2A-KO cell lines were kindly provided by Dr. Andrew P. Jackson’s lab to detect slippage mutation rate at STRs. We further knocked out the PARP1 gene in these three cell lines (see “Cell Lines” in this methods section). The fluctuation assays were performed as described by Dr. Andrew P. Jackson’s team.^12^ For the PARP inhibitor pre-treatment in the fluctuation assay, we treated our cells in a flask with either 1 μM talazoparib or a similar volume of DMSO (for the negative control) for 3 days before plating them for the fluctuation assay as outlined in the paper by Reijns, *et al*.

### Cell proliferation analysis

To assess the proliferation of the different strains, 20,000 cells were seeded in multiple 12-well plates (with triplicates in each plate, and each plate representing a different day to be read, from day 0 to 5). Six hours after plating the day 0 plate, the plate was washed with 1x PBS. After washing, the plates were fixed with a buffer consisting of 1% formaldehyde, 0.05% Crystal violet (w/v), 1x PBS, and 1% methanol for 20 mins. The wells were washed three times with water and allowed to air dry overnight. Once dried, 2 mL of 10% acetic acid was added to each well and incubated for 20 mins with moderate shaking. The wells were mixed with a pipette, and the plates were measured for 590 nm absorbance. Each day at the same time, a plate was fixed and allowed to air dry. Once the data points for all the days were obtained, the growth curve was plotted by normalizing the value of the signals using day 0 as the frame of reference.

### MTT viability assay

The MTT assay to assess the viability of cells after treatments has been described previously.^59^ To assess viability after treatments with different inhibitors, we plated the different strains each with 8 repeats (one full column) in a 96-well plate (1000-2000 cells per well). The following day, we treated them with the appropriate drug to be tested (untreated controls were also maintained for each strain). After the treatment period (6 days), each well was supplemented with 20 μL 5mg/mL thiazolyl blue tetrazolium bromide (MTT) solution (in PBS). The cells were then incubated for 3.5 hours in the 37 °C tissue culture incubator. The media was then removed and replaced with 150 μL MTT solvent (4 mM HCl and 0.1% NP-40 diluted in isopropanol), covered with aluminum foil, and incubated in a shaker for 15 minutes. The absorbance at 590 nM with a 620 nM reference for all the wells was measured using an Epoch plate reader (Biotek). The results were normalized to the appropriate untreated controls for each strain.

### TOP1cc quantification RADAR assay

The genomic DNA purification was conducted using the TOPOGEN ICE Assay Kit (TG10201),^35,36^ except that the cells were treated with 10 μM MG-132 protease inhibitor for 2 hours before DNA extraction to protect the TOP1-DNA adducts from degradation. Cells pre-treated with 20 μM camptothecin for 2-hour pre-DNA extraction were included as a positive control. The concentration of genomic DNA was quantified by nanodrop before and after dilution to 30 ng/μL using the sodium phosphate buffer. Each sample, 200 μL, was vacuum-blotted into both a nitrocellulose membrane, for protein detection, and an Amersham Hybond -N^+^ membrane (GE Healthcare), for DNA detection (which also serves as a loading control). After blotting, both membranes were irradiated with 120 mJ/cm^2^ using a UV Stratalinker 1800 (Stratagene). The nitrocellulose membrane was washed with TBST and immuno-stained with TOP1 antibody (provided in the TOPOGEN ICE kit) in 5% skim milk (w/v), TBS with 0.1% Tween-20 (v/v), while the Hybond membrane was washed with TBS and stained with Sybr Gold (Invitrogen) diluted in TBS. For detection of the TOP1 immuno-stained membrane, we used IRDye 800CW goat anti-rabbit antibody (LICOR). They were both imaged using a ChemiDoc MP imaging system (BIO-RAD).

### Co-immunoprecipitation assay

U2OS cells were generated expressing either the wild-type or the R735E mutant of a 3xFLAG-tagged PARP1 in the PARP1-KO U2OS cells (a kind gift from Dr. Shan Zha).^60^ The pCMV-PARP1-3xFlag-WT plasmid was a gift from Thomas Muir’s lab (Addgene #111575).^61^ To assess TOP1-PARP1 interaction in cells, 1 million cells were pelleted and frozen at -80 °C. The pellets were then resuspended in 400 μL PBS buffer supplemented with 0.01% NP-40, 1mM β-mercaptoethanol, and a protease inhibitors cocktail (aprotinin, chymostatin, leupeptin, and pepstatin A at 5 μg/ml each, and 1 mM phenyl-methylsulfonyl fluoride). The samples were sonicated at 40% amplification in a Fisher Scientific Model 120 sonicator for 5 secs to achieve proper extraction of the nuclear proteins. The lysate mixture was incubated with 25 μL anti-FLAG M2 agarose resin (Sigma) overnight at 4 °C. Following incubation, the resin was washed two times with 400 μl of wash buffer (1x PBS, 0.1% NP-40, 1mM β-mercaptoethanol) and eluted with 20 μl of 2x Laemmli sample buffer (BIO-RAD). The input and SDS eluate fractions were resolved by 10% SDS-PAGE and immunoblotted with either anti-TOP1 (Cell signaling # 79971S), anti-PARP1 (Millipore Sigma #SAB5701291), or anti-GAPDH (BioLegend #649203) antibody (the latter as a loading control). IRDye 800CW goat anti-rabbit antibody (LICOR # 926-32211) was used as secondary antibody for protein detection.

## Acknowledgements

We are grateful to Shan Zha, Andrew P. Jackson, John Pascal, Thomas Muir, and Joan Conaway for providing plasmids and human cell lines. Research reported in this publication was supported by the National Cancer Institute of the National Institutes of Health (NIH) under Award Numbers F31CA268751 (E.S.) and R01CA285684 (H.N. and Q.C.). Part of effort for H.N was also supported by R35GM152207 and the American Cancer Society Research Scholar Award RSG-21-013-01-DMC. Part of effort for Q.C. was also supported by R01CA256741, R01CA278832 and R01CA300246, Prostate SPORE P50CA180995 Development Research Program and Polsky Urologic Cancer Institute of the Robert H. Lurie Comprehensive Cancer Center of Northwestern University at Northwestern Memorial Hospital.

---

**Supplementary Table 1:** List of plasmids used.

| Construct | Description | reference |
| --- | --- | --- |
| pFASTBac-HTB- FLAG-His<sub>6</sub>-TOP1 | Plasmid used to produce the Bacmid containing the 6xHis/FLAG-tagged TOP1 gene for expression in insect cells. | This work |
| pFASTBac-HTB- FLAG-His<sub>6</sub>-TOP1-Y723F | Plasmid used to produce the Bacmid containing the His6/FLAG-tagged TOP1-Y723F gene for expression in insect cells. | This work |
| pET28a-His<sub>6</sub>-PARP1 | Plasmid for expressing full length PARP1 in *E. coli*, a kind gift from John Pascal's lab. | <sup>1</sup> |
| pET28a-His<sub>6</sub>-PARP1-K695E | Plasmid for expressing PARP1-K695E mutant in *E. coli*. | This work |
| pET28a-His<sub>6</sub>-PARP1-K700E+R704E | Plasmid for expressing PARP1-K700E+R704E mutant in *E. coli*. | This work |
| pET28a-His<sub>6</sub>-PARP1-R735E | Plasmid for expressing full length PARP1-R735E in *E. coli*. | This work |
| pETDuet-His<sub>6</sub>-PARP1-ΔDBD-FLAG | Bacterial expression of PARP1 lacking the DBD domain (379-1014). | This work |
| pETDuet- His<sub>6</sub>-PARP1-N-366 | Bacterial expression of DBD domain of PARP1. | This work |
| pET28_ His<sub>6</sub>-PARP1-ΔCAT (Addgene# 173942) | Bacterial expression of PARP1 lacking the CAT domain. | <sup>2</sup> |
| pET28_ His<sub>6</sub>-PARP1-ΔART (Addgene #174804) | Bacterial expression of an inactive PARP1 mutant lacking the ART subdomain. | <sup>2</sup> |
| pcDNA3.1/Zeo(+)-TDP1 | Plasmid for mammalian overexpression of TDP1. | This work |
| pCMV-PARP1-3xFlag (Addgene # 111575) | Plasmid for mammalian overexpression of 3xFLAG-PARP1 | <sup>3</sup> |
| pCMV-PARP1-R735E-3xFLAG | Plasmid for mammalian overexpression of 3xFLAG-PARP1-R735E mutant | This work |

**Supplementary Table 2:**
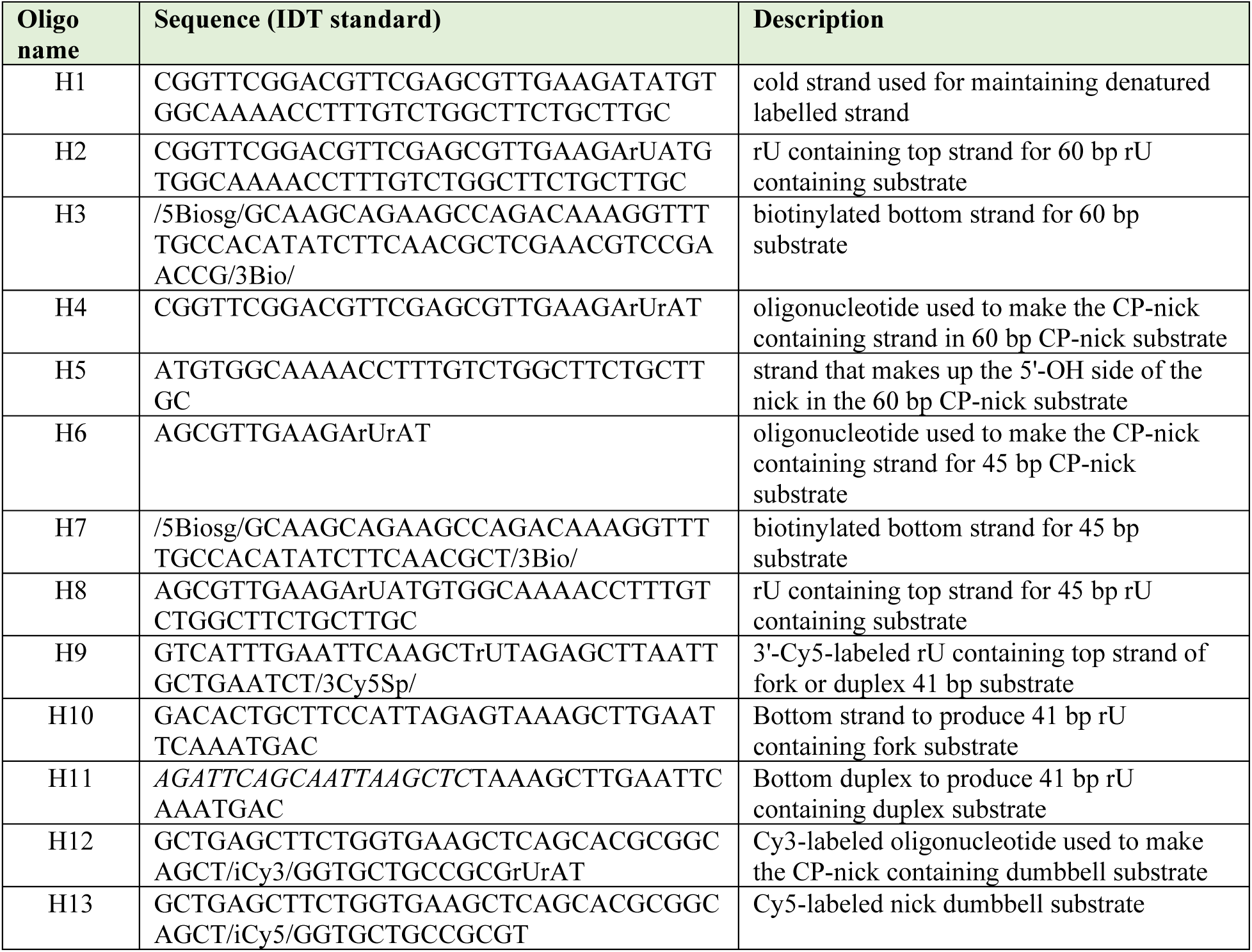
List of oligonucleotides used to produce the substrates for biochemical assays.

**Supplementary Table 3:**
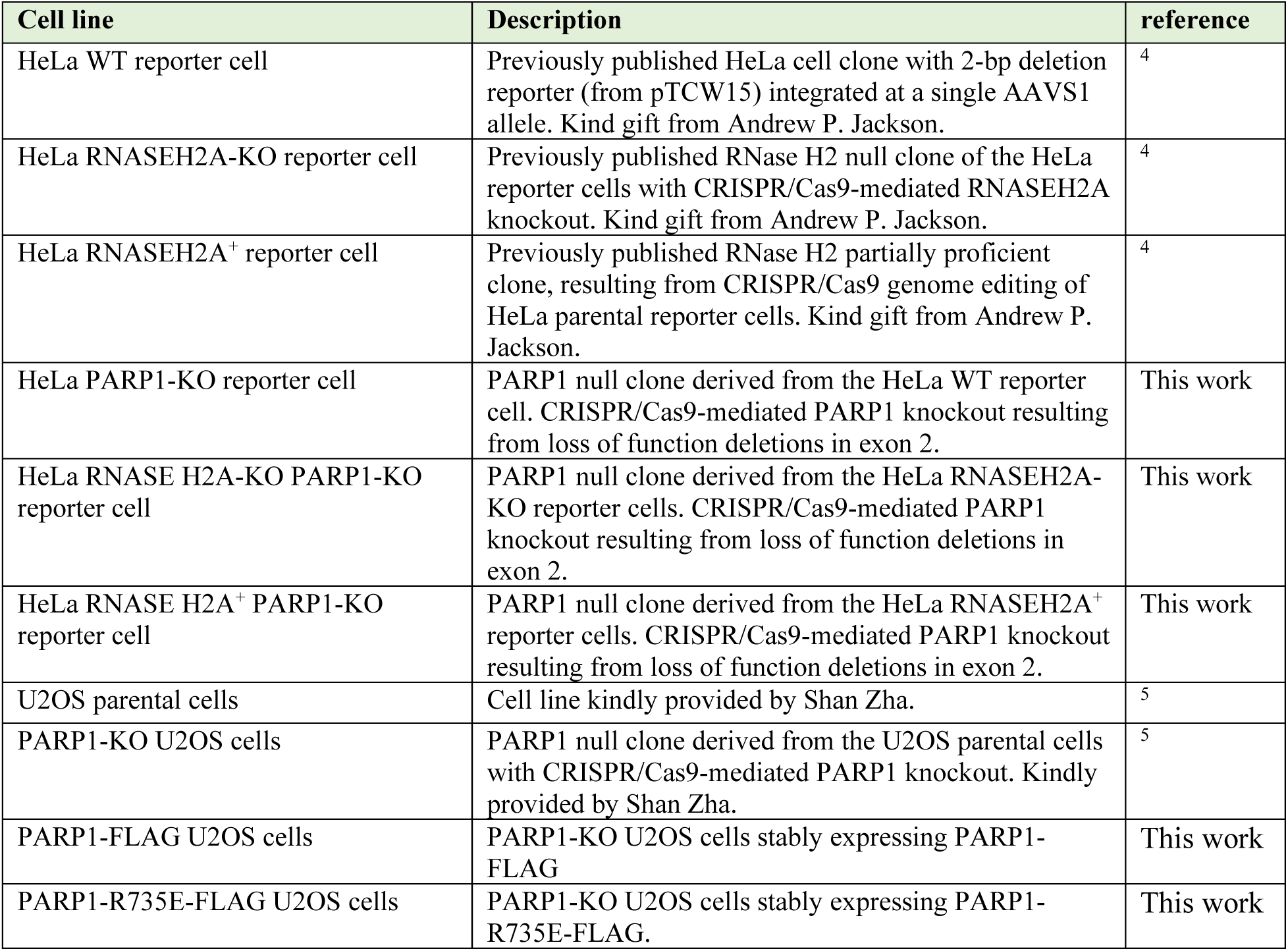
List of cell lines used.

| Cell line | Description | reference |
| --- | --- | --- |
| HeLa WT reporter cell | Previously published HeLa cell clone with 2-bp deletion reporter (from pTCW15) integrated at a single AAVS1 allele. Kind gift from Andrew P. Jackson. | <sup>4</sup> |
| HeLa RNASEH2A-KO reporter cell | Previously published RNase H2 null clone of the HeLa reporter cells with CRISPR/Cas9-mediated RNASEH2A knockout. Kind gift from Andrew P. Jackson. | <sup>4</sup> |
| HeLa RNASEH2A<sup>+</sup> reporter cell | Previously published RNase H2 partially proficient clone, resulting from CRISPR/Cas9 genome editing of HeLa parental reporter cells. Kind gift from Andrew P. Jackson. | <sup>4</sup> |
| HeLa PARP1-KO reporter cell | PARP1 null clone derived from the HeLa WT reporter cell. CRISPR/Cas9-mediated PARP1 knockout resulting from loss of function deletions in exon 2. | This work |
| HeLa RNASE H2A-KO PARP1-KO reporter cell | PARP1 null clone derived from the HeLa RNASEH2A-KO reporter cells. CRISPR/Cas9-mediated PARP1 knockout resulting from loss of function deletions in exon 2. | This work |
| HeLa RNASE H2A<sup>+</sup> PARP1-KO reporter cell | PARP1 null clone derived from the HeLa RNASEH2A<sup>+</sup> reporter cells. CRISPR/Cas9-mediated PARP1 knockout resulting from loss of function deletions in exon 2. | This work |
| U2OS parental cells | Cell line kindly provided by Shan Zha. | <sup>5</sup> |
| PARP1-KO U2OS cells | PARP1 null clone derived from the U2OS parental cells with CRISPR/Cas9-mediated PARP1 knockout. Kindly provided by Shan Zha. | <sup>5</sup> |
| PARP1-FLAG U2OS cells | PARP1-KO U2OS cells stably expressing PARP1-FLAG | This work |
| PARP1-R735E-FLAG U2OS cells | PARP1-KO U2OS cells stably expressing PARP1-R735E-FLAG. | This work |

**Supplementary Table 4:**
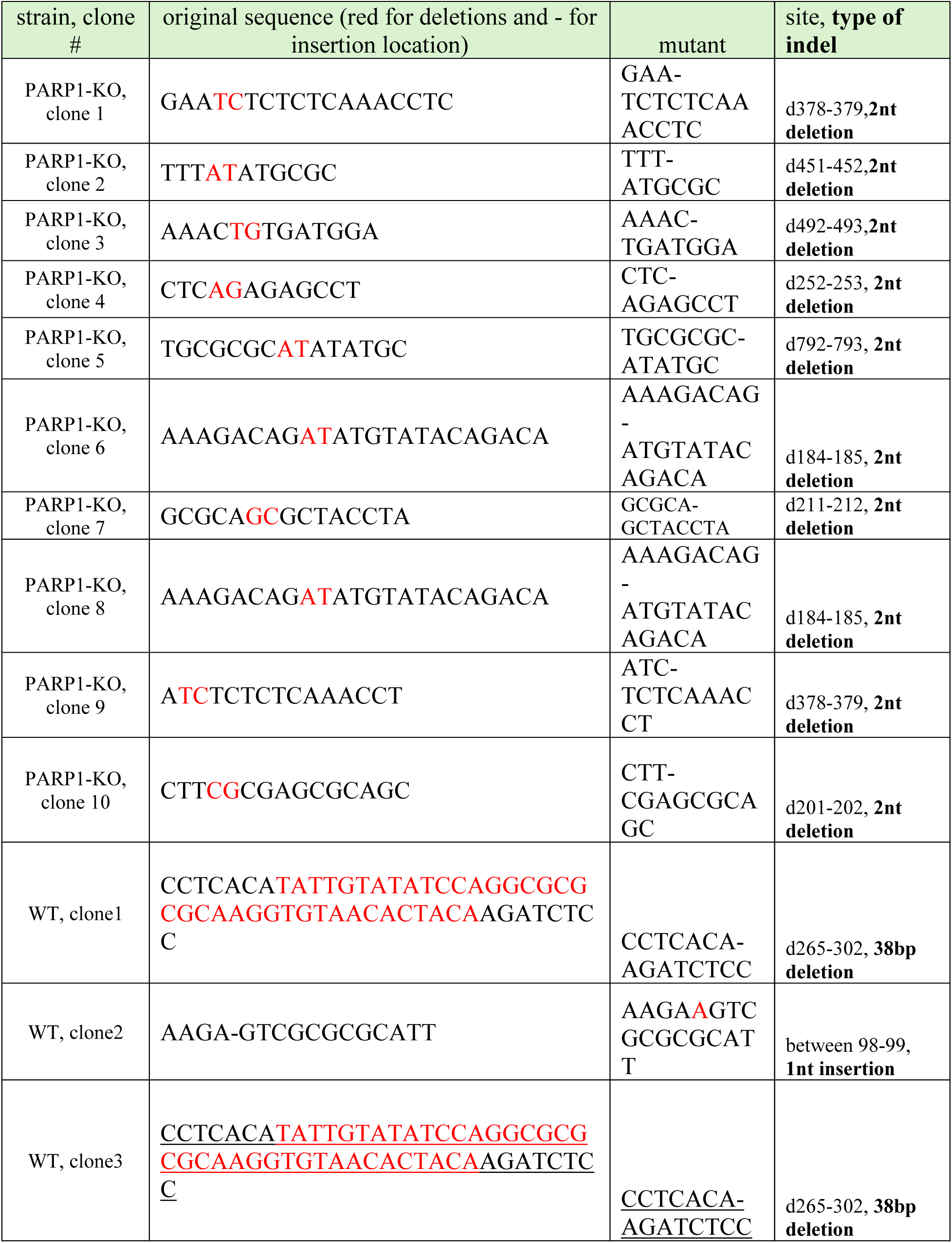
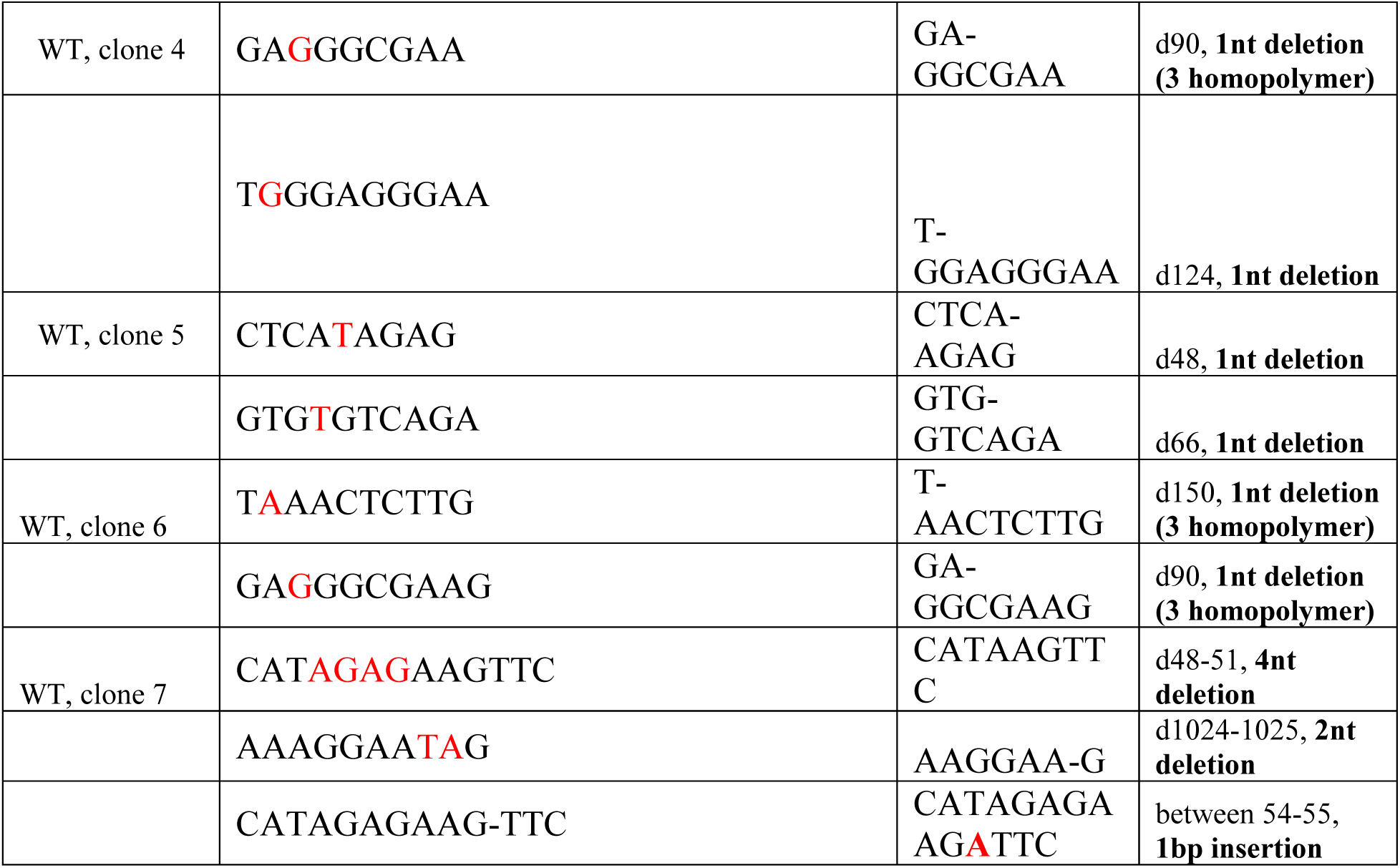
List of mutations found in the hygromycin-puromycin fusion reporter gene of isolated clones of either PARP1-KO or WT strains.

**Supplemental Figure 1:**
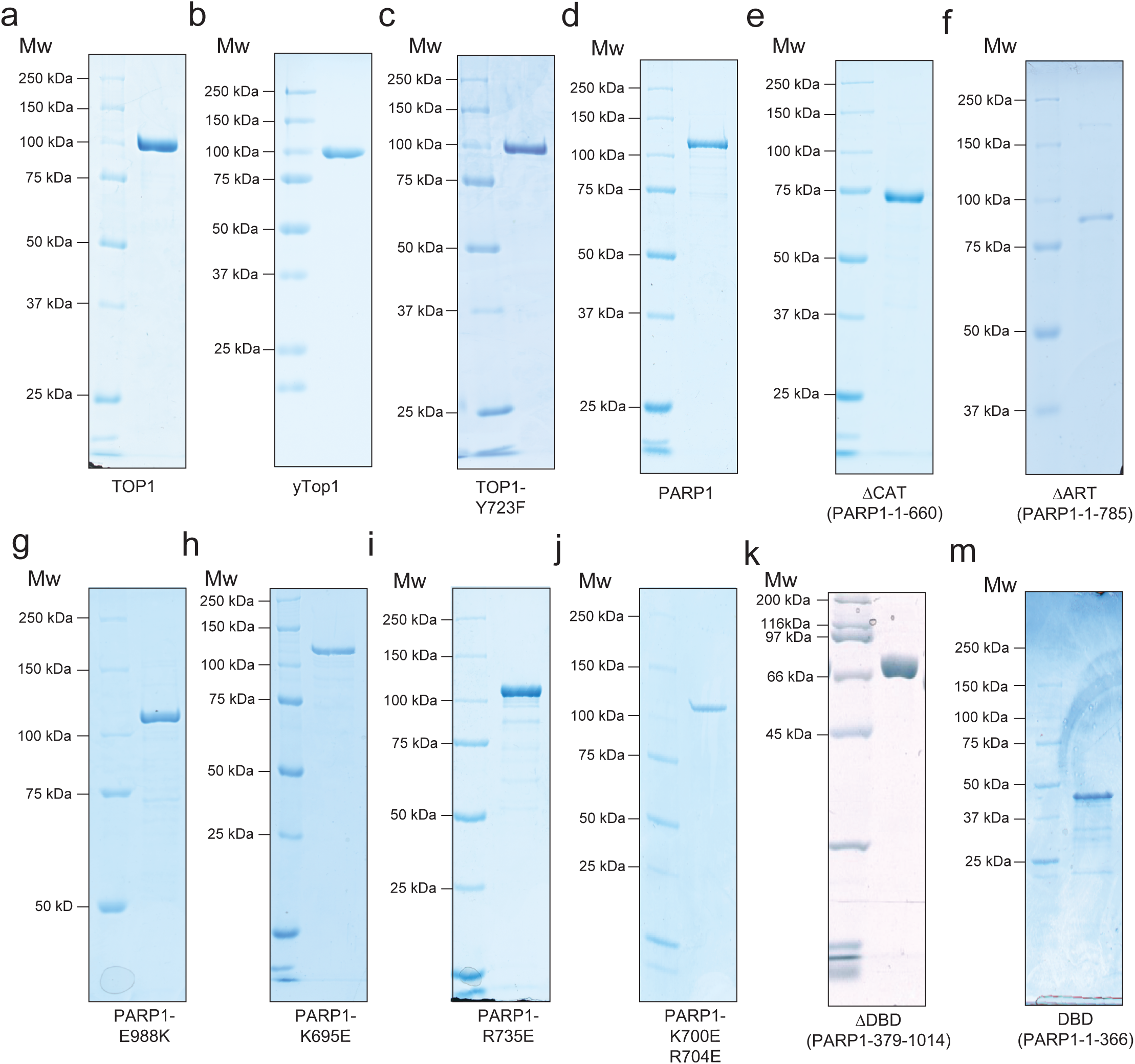
SDS-PAGE analysis of purified proteins. Purified proteins were fractionated in 4-15% SDS-PAGE or 10% SDS-PAGE and stained by Coomassie blue G-250.

**Supplemental Figure 2:**
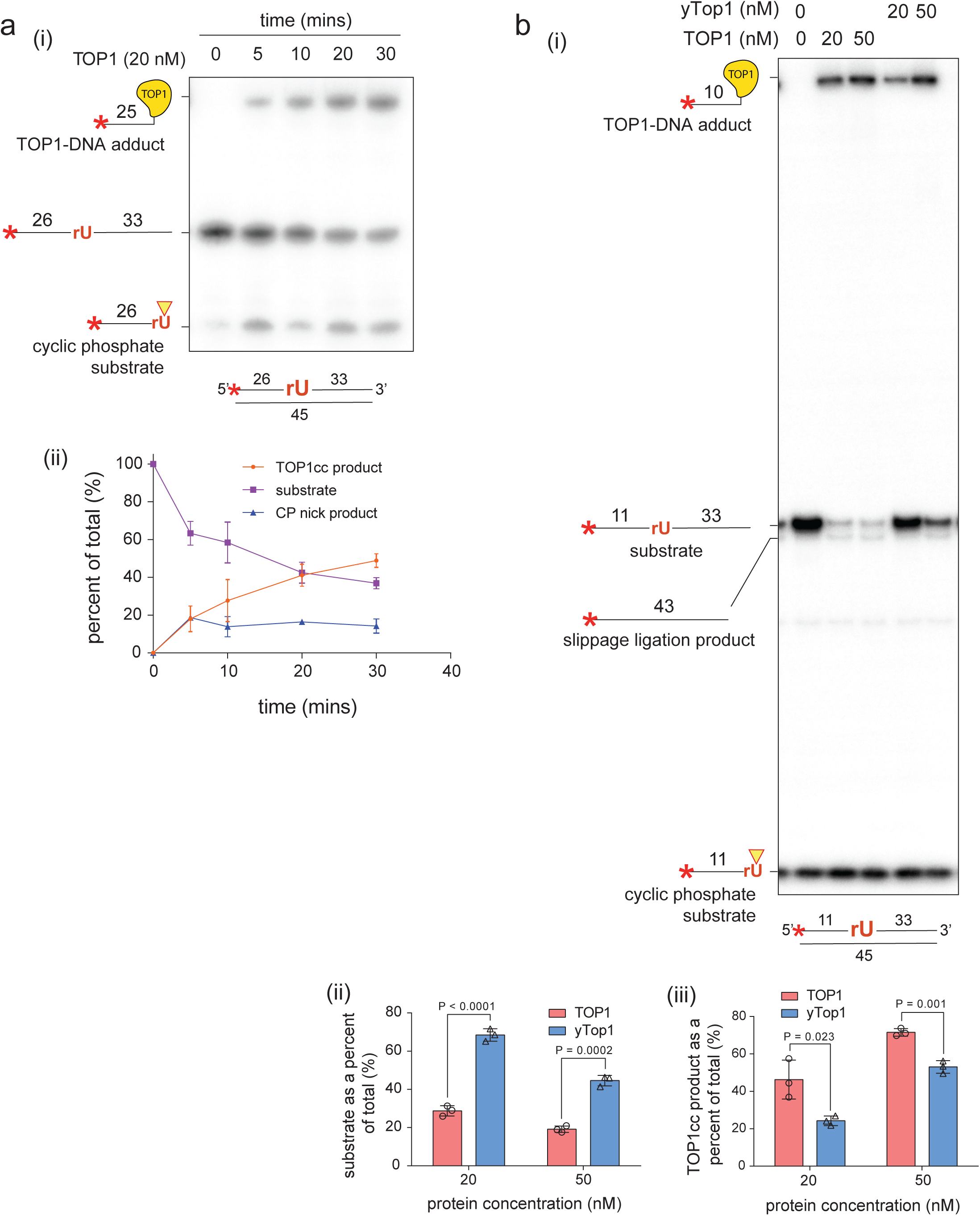
Human TOP1 rNMP cleavage analysis and comparison with yeast Top1 (yTop1). a) (i) Time course analysis of ribonucleotide processing by human TOP1. (ii) Quantification of different product species and the remaining substrate. b) (i) Comparison of ribonucleotide processing between human TOP1 and yTop1. (ii)-(iii) Quantification of the remaining substrate and the TOP1cc product for each TOP1 variant.

**Supplemental Figure 3.**
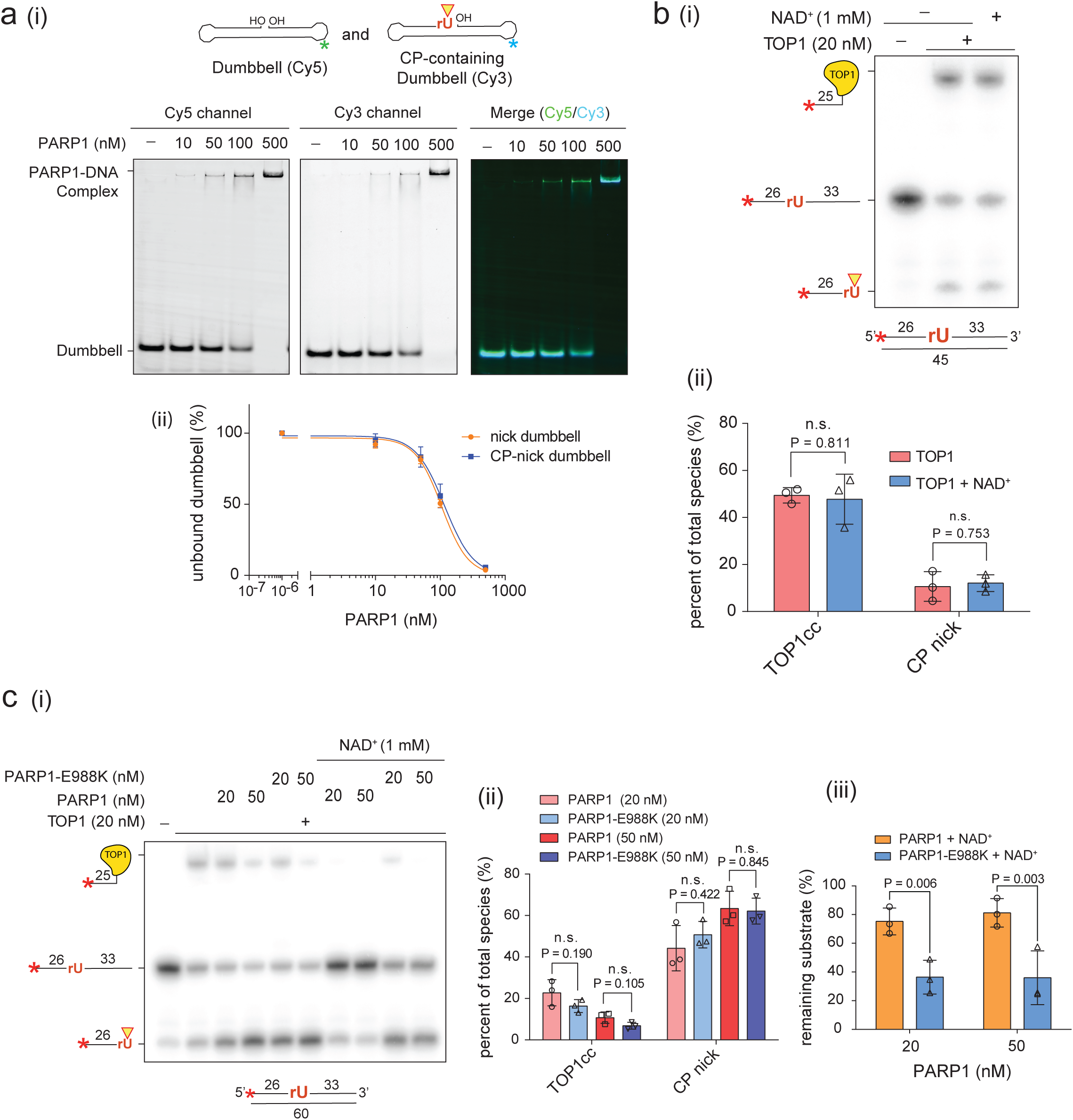
CP nick activates PARP1 for TOP1 regulation. a) (i) PARP1 binding competition assay using a Cy5 labeled dumbbell nick substrate and a Cy3 labeled dumbbell CP-nick substrate. Single channel and merged images are shown. (ii) Quantification of the remaining unbound substrate for each condition. b) (i) TOP1 ribonucleotide cleavage assay performed in the absence and presence of NAD^+^ to examine the effect of NAD^+^ on TOP1. (ii) quantification of TOP1cc and CP nick products. c) (i) TOP1 ribonucleotide cleavage assay performed in the absence or presence of indicated amount of PARP1 or the catalytically dead PARP1-E988K mutant, with or without NAD^+^. (ii) Quantification of TOP1cc and CP nick products from each reaction. (iii) Quantification of the remaining substrate in reactions containing PARP1 or PARP1-E988K in the presence of NAD^+^.

**Supplemental Figure 4:**
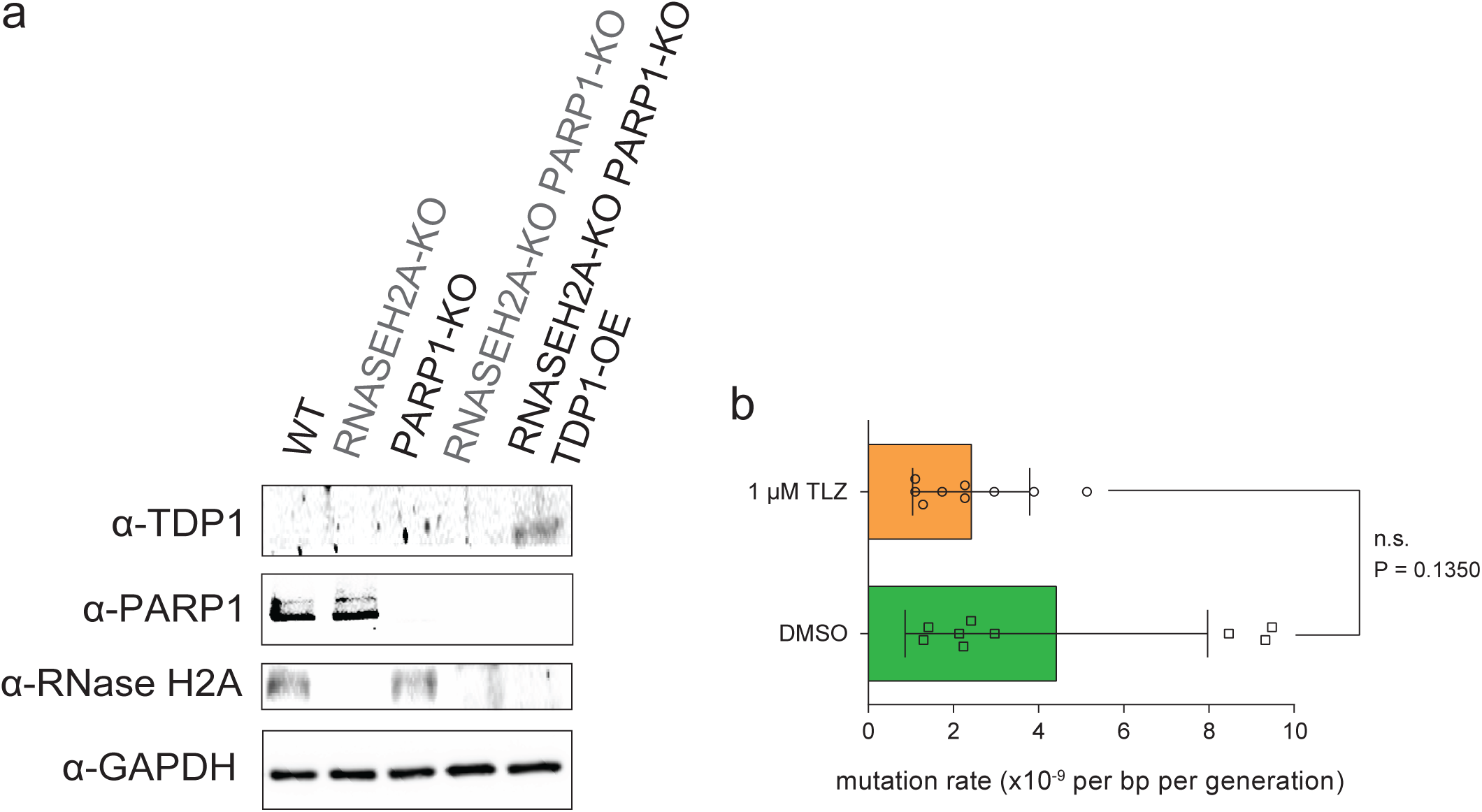
Confirmation of PARP1 knockout and TDP1 overexpression in the reporter cell lines and examination of the effect of PARP inhibitor on the mutation rate in the wild-type reporter cells. a) PARP1 was knocked out in the WT, RNase H2A-KO, and RNase H2A+ reporter HeLa cells. TDP1 was stably expressed in the RNase H2A-KO PARP1-KO cell line under the control of cmv promoter. The protein samples were fractionated in 4-15% SDS-PAGE and transferred into a nitrocellulose membrane for western blots. b) Fluctuation assay reporting on the mutation rate of the WT HeLa reporter strain pre-treated with either PARP inhibitor talazoparib (TLZ) or DMSO, showcasing that inhibition of PARP1 does not significantly affect the number of deletions at STRs.

**Supplementary Figure 5:**
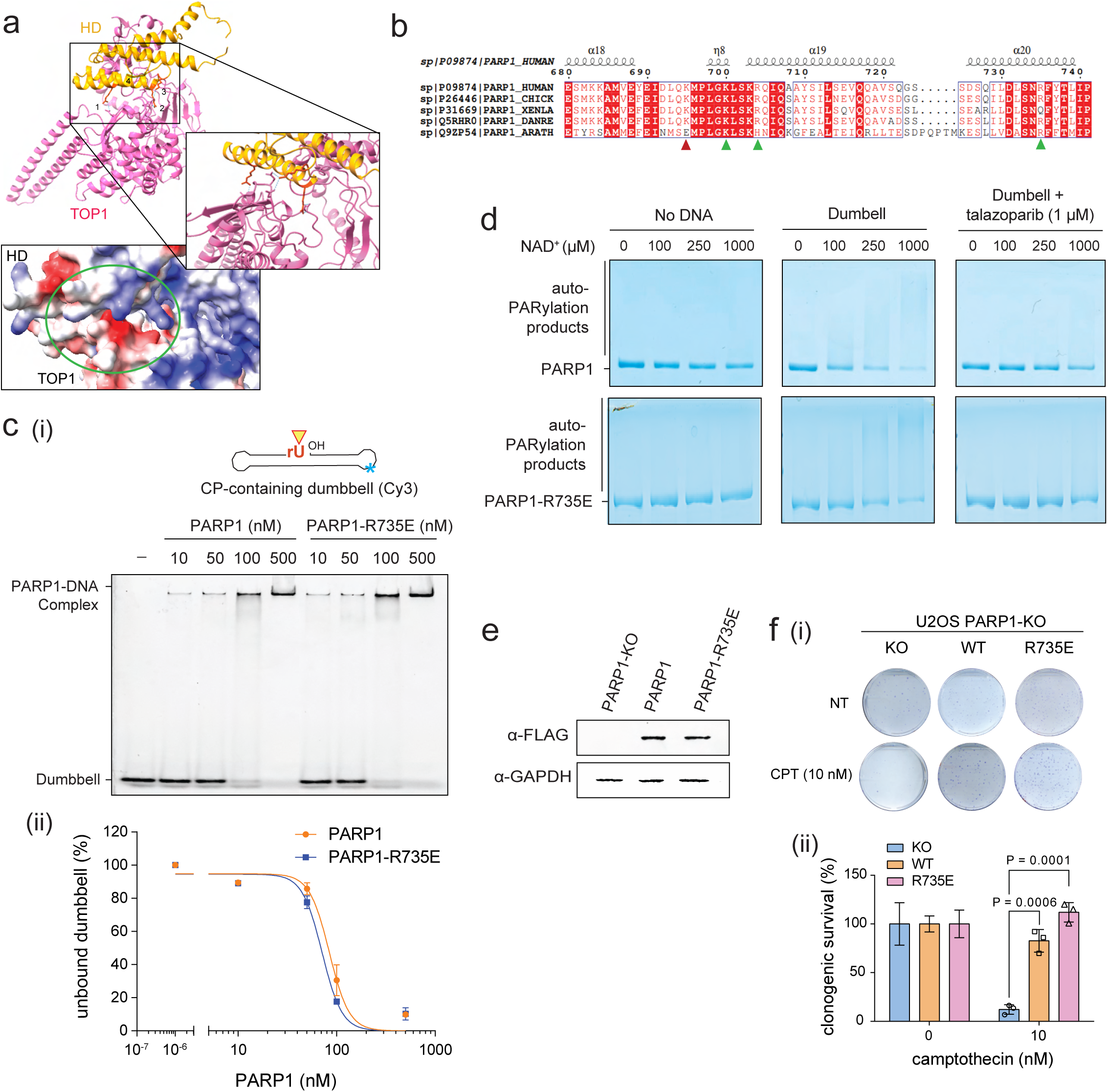
Though deficient in TOP1-interaction, PARP1-R735E is comparable to the wild-type in DNA binding, autoPARylation and response to PARP inhibitor *in vitro*, as well as complementing PARP1-KO for the sensitivity to camptothecin in cells. a) A model of the interface between TOP1 and the HD domain of PARP1 from AlphaFold prediction. The targeted residues within the the HD domain (K695, K700, R704, and R735) are labeled with the numbers 1-4, respectively. A close-up of the PARP1-TOP1 interface and the electrostatic surface map of this interface. b) Alignment of the PARP1 HD domain region containing the targeted residues across higher eukaryotes, including human, chicken, Xenopus frog, zebrafish, and Arabidopsis thaliana. Arrows mark the sites tested for TOP1 interactions. Green arrows indicate sites whose mutation reduced TOP1 binding, whereas the red arrow indicates a site that retained substantial TOP1 binding after mutation. c) (i) DNA binding assay comparing WT PARP1 and the R735E mutant ability to bind to the CP-nicked dumbbell substrate. (ii) Quantification of the unbound substrate for both PARP1 variants. d) Coomassie stained 10% SDS-PAGE images showcasing the auto-PARylation activity and talazoparib sensitivity of both PARP1 and PARP1-R735E. e) Western blot showcasing the stable expression of PARP1-FLAG and PARP1-R735E-FLAG in the PARP1-KO U2OS cell lines. f) (i) Clonogenic assay assessing the sensitivity to camptothecin (CPT) of either the PARP1-KO U2OS cell line or the PARP1-KO cell line expressing either WT PARP1 or PARP1-R735E. (ii) Quantification of clonogenic survival for each strain.

